# Mutant IDH impairs chromatin binding by PDGFB to promote chromosome instability

**DOI:** 10.1101/2025.02.20.639365

**Authors:** Rachel N. Curry, Malcolm F. McDonald, Peihao He, Brittney Lozzi, Yeunjung Ko, Isabella O’Reilly, Anna Rosenbaum, Wookbong Kwon, Leyla Fahim, Joshua Marcus, Noah Powell, Su Wang, Jin Ma, Asha Multani, Dong-Joo Choi, Debo Sardar, Carrie Mohila, Jason Lee, Marco Gallo, Arif Harmanci, Akdes Serin Harmanci, Benjamin Deneen, Ganesh Rao

## Abstract

Non-canonical roles for growth factors in the nucleus have been previously described, but their mechanism of action and biological roles remain enigmatic. Platelet-derived growth factor B (PDGFB) can drive formation of low-grade glioma and here we show that it localizes to the nucleus of human glioma cells where it binds chromatin to preserve genome stability and cell lineage. Failure of PDGFB to localize to the nucleus leads to chromosomal abnormalities, aberrant heterochromatin architecture and accelerated tumorigenesis. Furthermore, nuclear localization of PDGFB is reliant upon the expression levels and mutation status of isocitrate dehydrogenase (IDH). Unexpectedly, we identified macrophages as the predominant source of PDGFB in human, finding that immune-derived PDGFB can localize to the nucleus of glioma cells. Collectively, these studies show that immune derived PDGFB enters the nucleus of glioma cells to maintain genomic stability, while identifying a new mechanism by which IDH mutations promote gliomagenesis.

## Introduction

Platelet-derived growth factor B (PDGFB) is a potent mitogen with established roles in both development and cancer^1–3^. In the central nervous system, PDGFB interacts with its receptor PDGFRA on oligodendrocyte precursor cells (OPCs) to regulate proliferation and maintain OPC lineage by preventing differentiation into oligodendrocytes (OLs)^4–6^. PDGFB binding to PDGFRA, a classical receptor tyrosine kinase (RTK), activates a series of phosphorylation events that culminate in changes to cell proliferation and cell states^7,8^. PDGFB is frequently used to induce glioma in mouse models and amplifications to PDGFRA are frequent in glioma^9^ and other cancers^10–12^. Although the mitogenic effects are attributed to their RTK activity, RTK-independent roles for PDGFRA have been demonstrated^13^. Among these, tumorigenic functions of PDGFRA variants in the cytoplasm of glioma cells^14^, along with nuclear roles for other RTKs^15^, provide evidence for non-canonical roles for growth factor receptors. These observations raise the question of whether growth factor ligands associated with RTKs can also localize to the nucleus. Indeed, nearly 40 years ago, prior studies demonstrated that exogenously-produced PDGFB binds to chromatin in cells that bear the cognate receptors^16^. Despite these precedents, whether growth factors exist in the nucleus remains unclear and their biological significance in the pathogenesis of glioma is unknown.

In adult glioma, mutations to isocitrate dehydrogenase (IDH) define a type of malignant brain tumors that has significantly longer survival outcomes than the IDH wild-type (wtIDH) counterparts^17,18^. In contrast to wtIDH glioma, which are attributed to driver mutations frequently occurring in cell cycle-related genes, IDH mutant (mIDH) gliomas feature mutations to one of two IDH genes, *IDH1* or *IDH2,* the majority of which are R132H and R172X, respectively^18^. It has been shown that canonical IDH mutations lead to neomorphic activity that yields overproduction of D-2-hydroxyglutarate (D2HG)^19^, a metabolite that has various oncogenic effects^20–23^. In mouse models, however, the canonical IDH1^R132H^ mutation is insufficient to drive gliomagenesis without secondary genomic drivers^24–26^, suggesting that other additional oncogenic events are required to induce tumors. Nevertheless, given their ubiquitous appearance in mIDH tumors, IDH mutations are thought to be an early transforming event in mIDH gliomagenesis. To date, however, a clear understanding of how IDH mutations mechanistically initiate glioma has yet to be achieved.

Like most other cancers, glioma is marked by genomic instability, which is defined as the propensity of the genome to incur small or large genomic aberrations^27^. Proper partitioning of the genome throughout the cell cycle is a tightly regulated process that serves as a critical component of maintaining genomic integrity. Infidelities in these processes are frequently encountered in cancer^28^ and on a large genomic scale manifest as chromosomal instability, which includes aneuploidy or chromosomal rearrangements that can be sufficient to initiate tumorigenesis^29–31^. Recent studies have demonstrated that the abundance of whole-arm chromosome aneuploidy and rearrangements in cancer may result from instability at constitutive heterochromatin regions like the centromere – the genomic locus at which the kinetochore and mitotic spindle attaches to separate sister chromatids and the site at which chromosome *p* and *q* arms conjoin^32–34^. Prior work has shown that centromeric instability and aberrant heterochromatin formation is sufficient to cause aneuploidy and infidelities in chromatid segregations^35,36^, suggesting that centromere aberrations could serve as initiating events in cancer. In the mIDH glioma subtype of oligodendroglioma, chromosomal instability frequently manifests as whole-arm codeletions of chromosomes *1p* and *19q*, which are a defining cytogenetic feature of the oligodendroglioma subtype. To date, however, whether improper heterochromatin maintenance at centromeric regions is sufficient to trigger genomic instability in glioma remains undefined.

We hypothesized that PDGFB and mutant IDH, which occur uniformly in mIDH glioma, may interact to promote glioma formation. To test this hypothesis, we utilized surgically-resected human glioma samples and developed a new immunocompetent *in utero* electroporation model of PDGFB-driven glioma to investigate a non-canonical role for PDGFB exists in the nucleus. Our studies suggest that a paracrine signaling loop exists in which mutant IDH inhibits chromatin binding of immune-derived PDGFB in glioma cells. Mechanistically, we find that PDGFB binds heterochromatin to maintain genome stability and that mutant IDH likely impedes PDGFB binding at the centromere. These data implicate aberrant interactions between mutant IDH and nuclear PDGFB as putative initiating events in mIDH glioma.

### PDGFB binds chromatin in glioma

To gain a better understanding of the role of PDGFB in human glioma, we began with an examination of how PDGFB expression correlates with clinical outcome. We performed Kaplan-Meier survival analysis of human glioma samples from The Cancer Genome Atlas (TCGA) and found that high expression of PDGFB independently predicts worse survival outcomes in mIDH glioma but has no effect on overall survival in IDH wild-type (wtIDH) glioma (**Figs. S1a–b**). In glioma, PDGFB and its cognate receptor, PDGFRA, are detected in mIDH tumors (**Fig. S1c**), whereas, in the non-tumor human brain, PDGFB is minimally expressed by microglia and vascular cells, and PDGFRA is highly expressed by OPCs (**Figs. S1d**). Given its correlation with poor survival outcomes and high expression levels in mIDH glioma, we sought to characterize PDGFB protein expression in mIDH tumors through immunostaining. We used antibodies specific for the canonical IDH1^R132H^ mutation and PDGFB to investigate the localization and distribution of PDGFB in mIDH glioma samples. Surprisingly, we found punctate PDGFB foci colocalizing with DAPI-stained DNA in the nucleus of approximately half of the cells in human samples (**Fig. 1a**). This nuclear PDGFB staining was observed in both IDH1^R132H–^ and IDH1^R132H^+ cells and appeared to localize near DAPI-rich heterochromatin, which is tightly packed regions of DNA commonly found near centromeres (**Fig. 1a**). These initial observations demonstrated that PDGFB expression is correlated with survival outcomes in mIDH glioma patients and is detected in the nuclei of cells in mIDH glioma.

**Figure 1.**
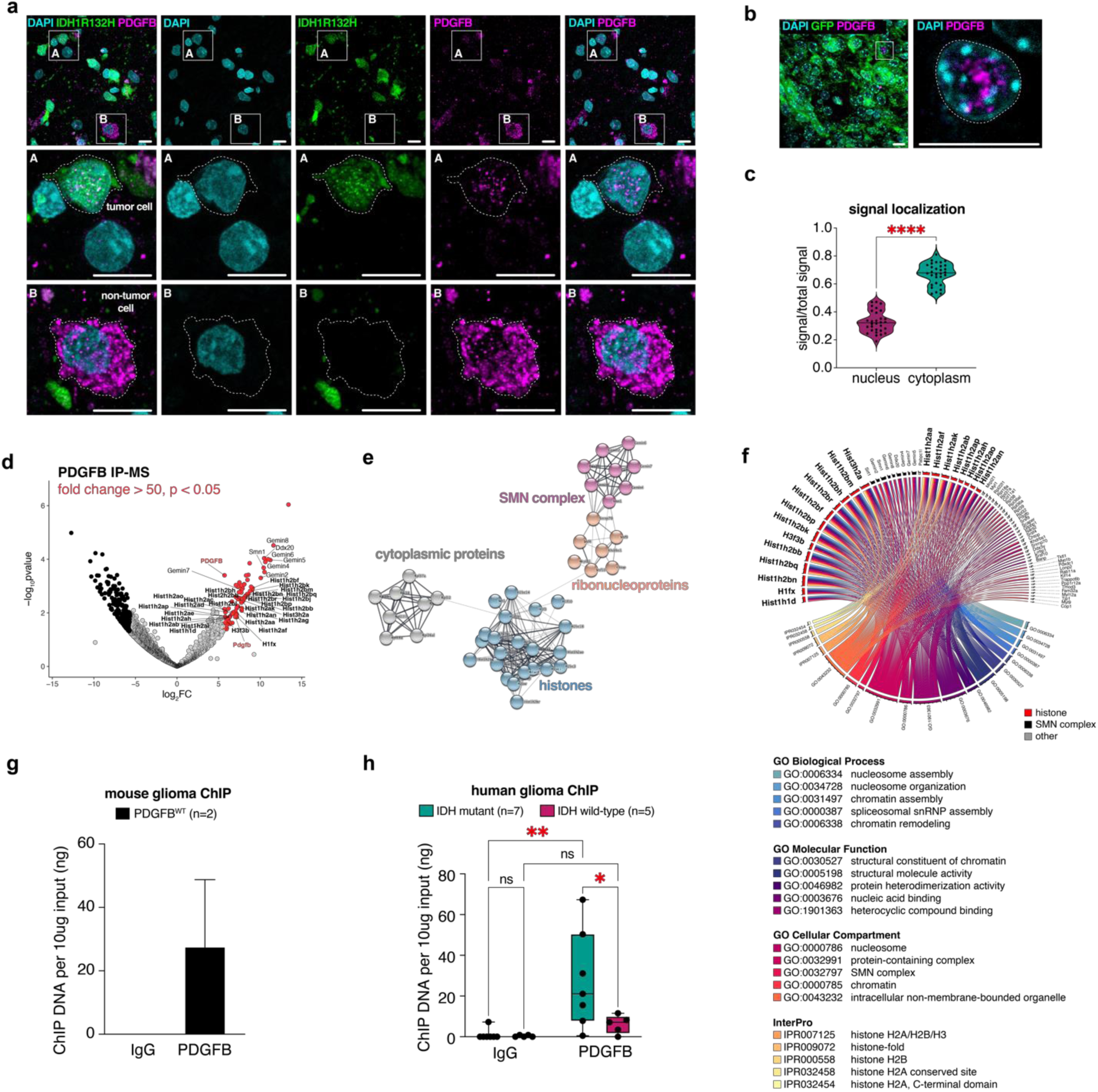
PDGFB binds chromatin in glioma. (a) Immunostaining of a surgically-resected human mIDH tumor sample shows PDGFB is detected in the nucleus in IDH1^R132H^+ tumor cells (inset A: white dashed line) and is detected in the cytoplasm and nucleus of IDH1^R132H^– non-tumor cells (inset B: white dashed line); scale bar = 20 microns. (b) Immunostaining of end-stage PDGFB^wt^ mouse brains shows PDGFB is found in the nucleus in GFP+ tumor cells, scale bar = 10 microns. (c) Quantification of immunofluorescence signal detected in the nucleus relative to total signal and cytoplasm relative to total signal in end-stage PDGFB^wt^ mouse brains; \*\*\**p*<0.0001. (d) Volcano plot of the 88 proteins identified from PDGFB IP-MS of PDGFB^wt^ tumors. Red dots denote proteins with fold change>50; *p*<0.05; black dots denote proteins enriched in IgG fold change<50; *p*<0.05. (e) STRING network of significant proteins from *(d)* showing enrichment of histones in PDGFB IP-MS. (f) Circos plot of GOs (*p*<0.05) corresponding to significant proteins from *(d).* (g) Bar graph of ChIP DNA shows PDGFB pulls down chromatin in PDGFB^wt^ mouse brains. (h) Bar graph of ChIP DNA shows PDGFB pulls down chromatin in human mIDH glioma (n=7; \*\**p*<0.01) but does not in human wtIDH glioma (n=5; ns). IP-MS: immunoprecipitation mass spectrometry; mIDH: IDH mutant.

To study the role of nuclear PDGFB in glioma progression, we developed a new model of immunocompetent glioma using *in utero* electroporation (IUE) of piggyBac transposase driving GFP and PDGFB overexpression (PDGFB^wt^) in neural progenitor cells (**Figs. S1e–f**). This model successfully recapitulates mIDH glioma disease course by generating *de novo* glioma in the absence of mutations. PDGFB^wt^ IUE mice have a median survival of 110 days, exhibit a low-to high-grade glioma (HGG) progression, and uniformly reproduce histopathological features of HGG including chromosome aberrations in the form of copy number variants (CNVs) (**Figs. S1g– j**). Similar to our analyses of human tumors, immunostaining of PDGFB^wt^ mice revealed analogous nuclear localization of PDGFB in tumor cells, which represents approximately 30% of PDGFB+ fluorescent signal, making the PDGFB^wt^ IUE system an optimal paradigm through which mechanistic studies of PDGFB can be executed (**Figs. 1b–c**). To discern the protein binding partners of PDGFB, we next performed PDGFB immunoprecipitation mass spectrometry (IP-MS) of end-stage PDGFB^wt^ tumors. Of the 88 proteins identified from IP-MS (fold change>50; *p*<0.05), 29 were histones (**Figs. 1d–e**; **Supplemental Table S1**). Corresponding gene ontology (GO) analysis of the 88 proteins revealed enrichment for nucleosome, chromatin and histone GOs (**Fig. 1f**). To determine if PDGFB binds DNA, we performed chromatin immunoprecipitation (ChIP) on PDGFB^wt^ induced mouse tumors (n=2) and surgically-resected human glioma samples (mIDH: n=7; wtIDH: n=5). The results of these experiments confirmed that PDGFB binds chromatin in mIDH glioma, which is paralleled in our model of PDGFB^wt^ mouse glioma (**Figs. 1g–h**). Collectively, these studies identify PDGFB as a novel chromatin-binding protein in mIDH glioma and describe a new immunocompetent model of low- to high-grade glioma.

### The C–terminal of PDGFB contains a highly conserved nuclear localization sequence

To understand the ability of PDGFB to localize to the nucleus, we investigated its nuclear localization sequence. PDGFB is produced as dimers of a 27.3kD preprotein that undergoes stepwise proteolytic processing at its N- and C-termini^37,38^. Cleavage occurs first at the N-terminal and yields an 18.4kD form of PDGFB that retains its C-terminus. Subsequent proteolysis occurs at the C-terminal to produce the mature 12.8kD form of PDGFB, which is considered the biologically active form of the protein (**Fig. 2a**). Surprisingly, our immunoblotting analyses revealed that the mature form of PDGFB was largely absent from mouse cortex, PDGFB^wt^ mouse glioma samples and human glioma samples (**Fig. 2b**). Instead, monomers or oligomers of either whole PDGFB preprotein (pre-PDGFB: 27.3kD), PDGFB preprotein without the N-terminal signal sequence (pre-PDGFB*; 24.9kD) or partially cleaved C-terminal-containing PDGFB (PDGFB-C; 18.4kD) were consistently found as the predominant forms of PDGFB in both murine and human brain whole cell lysates. Closer examination of the mature and propetide amino acid sequences of PDGFB revealed a high concentration of positively-charged basic residues at the C-terminal and that this C-terminal propetide was strongly predicted to bind DNA^39^ (**Figs. S2a**). This positively-charged amino acid compositional bias was also found to contain a putative nuclear localization sequence (NLS), which we identified using the structural motif-detecting algorithm, Eukaryotic Linear Motif (ELM)^40^. The NLS is highly conserved across more than 400 jawed vertebrate species and is more frequently mutated than any other region of PDGFB in cancer (**Figs. S2b–d**). Taken together, these data implicate the C-terminal as a biologically relevant component of PDGFB and suggest that the C-terminal bearing forms of pro-PDGFB and PDGFB-C are the predominant variants encountered in the mammalian brain.

**Figure 2.**
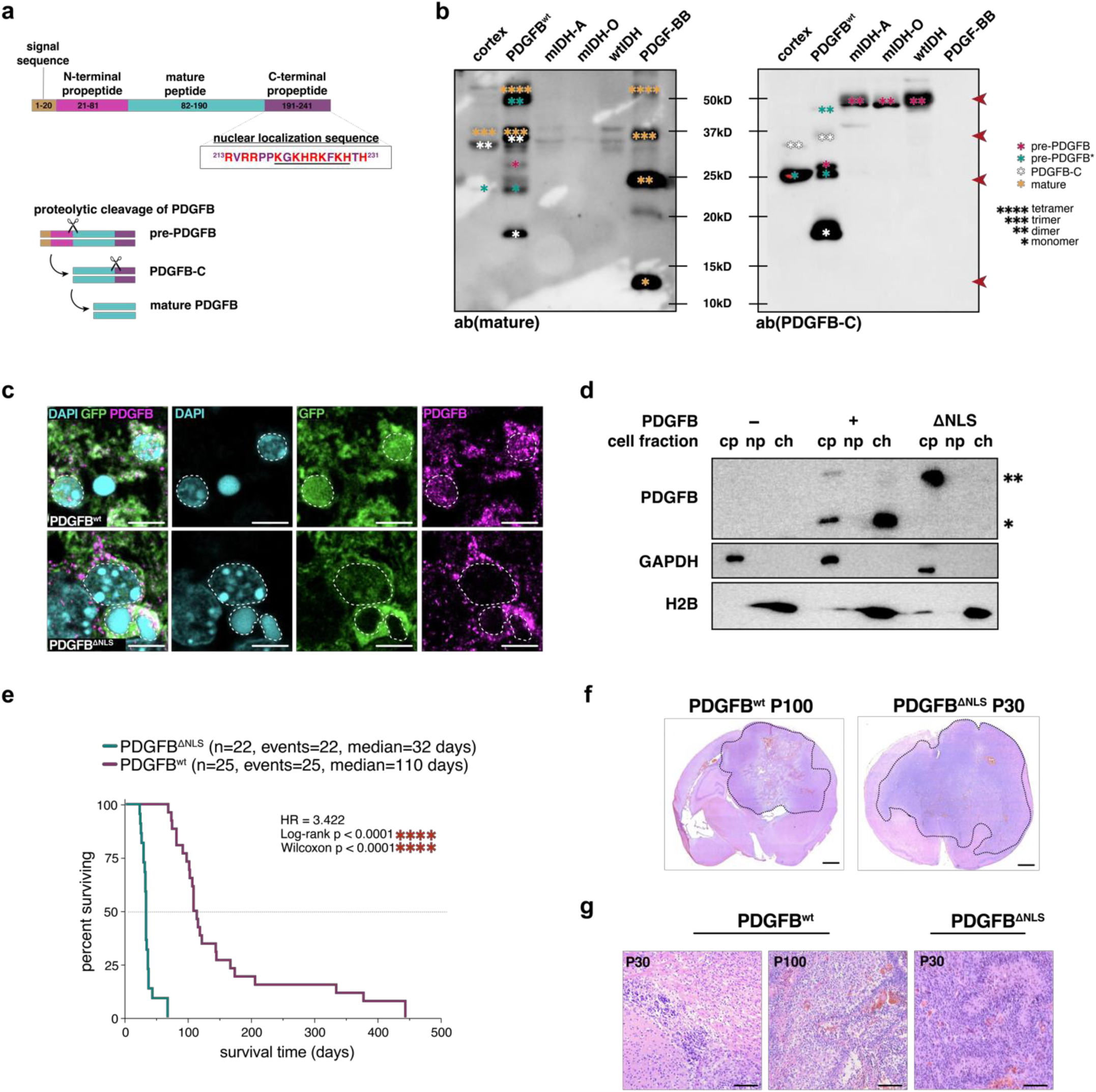
PDGFB contains a highly conserved nuclear localization sequence at its C–terminus. (a) Schematic representation of the step-wise proteolytic cleavage of PDGFB homodimers, PDGF-BB. Cleavage occurs at the N-terminus first and C-terminus secondarily. Underlined sequence denotes NLS. (b) Immunoblotting of mouse lysates (non-tumor cortex and PDGFB^wt^ tumor) and human glioma lysates. Immunoblotting using an antibody against mature PDGFB (left blot) shows mature PDGFB monomer (12.8kD) is not detected in mouse or human brain samples but is detected in recombinant PDGF-BB in monomeric (*), dimeric (**), trimeric (***) and tetrameric (****) forms. Immunoblotting using an antibody against the C-terminus of PDGFB (right blot) shows C-terminal-containing forms of PDGFB (PDGFB-C; 18.4kD), full pre-protein (pre-PDGFB; 27.3kD) and pre-protein without the signal sequence (pre-PDGFB*; 24.9kD) are detected in monomeric (*) and dimeric (**) forms in mouse and human brain samples. (c) Immunostaining of end-stage PDGFB^wt^ and PDGFB^ΔNLS^ mouse brains shows PDGFB is found in the nucleus in GFP+ tumor cells in PDGFB^wt^ tumors but is no longer detected in the nucleus of PDGFB^ΔNLS^ tumors; scale bar = 5 microns. (d) Immunoblots of cell fractions isolated from patient-derived wtIDH GSCs (7-2). PDGFB is present in both the cytoplasmic (cy) and chromatin (ch) fractions and is not present in the nucleoplasm (np) when PDGFB^wt^ is overexpressed. PDGFB is not detected in nuclear compartments when PDGFB^ΔNLS^ is overexpressed; * = 18.4kD; ** = 27.3kD; ch: chromatin fraction; np: nucleoplasm; cy: cytoplasmic fraction. (e) Kaplan-Meier survival analysis of PDGFB^wt^ (n=25, median=110 days) and PDGFB^ΔNLS^ (n=22; median=32 days) mice shows PDGFB^ΔNLS^ mice have significantly reduced survival times (HR=3.422; log-rank \*\*\*\**p*<0.0001; Wilcoxon *\*\*\*\*p*<0.0001). (f) H&E staining of a coronal section of end-stage PDGFB^wt^ (P100) and PDGFB^ΔNLS^ (P30) tumors. Black dashed line denotes tumor; scale bar = 2 mm. (g) H&E staining of PDGFB^wt^ and PDGFB^ΔNLS^ showing PDGFB^wt^ brains are histopathologically consistent with LGG at P30 and HGG at P100 whereas PDGFB^ΔNLS^ tumors are HGG at P30; scale bar = 100 microns. GSC: glioma stem cell; IP-MS: immunoprecipitation mass spectrometry; mIDH-A: astrocytoma; mIDH-O: oligodendroglioma; NLS: nuclear localization sequence.

To test whether the NLS sequence causes PDGFB nuclear localization, we neutralized the positively-charged arginine (R) and lysine (K) residues of PDGFB between amino acids 214-227 by converting them to serine (S); histidine (H), while also positively charged, confers important structural contributions to protein conformation and was therefore left unaltered to prevent unwanted secondary effects (**Fig. S2e**). We generated IUE mice using NLS-mutant PDGFB (PDGFB^ΔNLS^) and repeated IP-MS from microdissected GFP+ brain tissue, finding that PDGFB^ΔNLS^ no longer bound histones (**Figs. S2f–h**). Importantly, the other protein interactors previously identified from PDGFB^wt^ IP-MS were still present, suggesting that alterations to the K and R residues within the NLS uniquely impacted PDGFB-histone interactions. Consistent with these IP-MS data, immunostaining of PDGFB^ΔNLS^ confirmed that PDGFB was no longer detected in the nuclei of NLS-mutant mice, suggesting that PDGFB no longer bound histones due to an inability to translocate to the nucleus (**Fig. 2c**). To validate these findings in the context of human glioma, we obtained primary wtIDH glioma stem cells (GSCs), overexpressed PDGFB variants and performed immunoblotting of isolated cell fractions. Results demonstrated that PDGFB^wt^ is present in the chromatin fraction whereas PDGFB^ΔNLS^ is predominately found in the cytoplasm, confirming the NLS serves as a critical determinate of PDGFB nuclear localization (**Fig. 2d**). Kaplan-Meier survival analyses of PDGFB^ΔNLS^ mice showed a dramatic reduction in survival time with a median survival of 32 days as compared to 110 days in PDGFB^wt^ mice (**Fig. 2e**). End-stage brains harvested from both PDGFB^ΔNLS^ (P30) or PDGFB^wt^ (P100) mice displayed histopathological features consistent with HGG, whereas PDGFB^wt^ tumors at P30 lacked HGG hallmarks and were consistent with low grade glioma (**Figs. 2f–g**). Immunostaining confirmed that both tumor types were OLIG2+, suggesting that tumors formed from aberrant expansion of PDGFRA+ OPCs (**Fig. S3a**). *In vivo* and *in vitro* proliferation analyses of end-stage mice revealed that both tumor types were highly proliferative with PDGFB^ΔNLS^-overexpressing mIDH human glioma cells showing marginally increased proliferation over PDGFB^wt^ *in vitro* (**Figs. S3b–d**). In sum, these experiments confirm that the C-terminus of PDGFB contains a highly conserved NLS and that impediments to the ability of PDGFB to bind chromatin accelerates tumorigenesis.

### Nuclear PDGFB supports the OPC lineage

To elucidate how the presence of PDGFB in the nucleus impacts the chromatin landscape, we performed single cell RNA sequencing (scRNA-seq) and single cell transposase-accessible chromatin with sequencing (scATAC-seq) on PDGFB^ΔNLS^, PDGFB^wt^ and non-tumor mouse brains (**Figs. S3e–f**). Analysis of scRNA-seq confirmed that PDGFB^ΔNLS^ and PDGFB^wt^ transcripts colocalized with GFP in cells that were annotated by our SCRAM annotation pipeline^41^ as OPCs, cycling OPCs or oligodendrocytes (OLs) (**Figs. S4a–c**). Examining the proportions of cell types within each tumor revealed enrichments of OPCs and OLs in PDGFB^wt^ and PDGFB^ΔNLS^ tumors, respectively, implicating nuclear PDGFB in the differentiation of OPCs to OLs (**Figs. 3a–b**; **Supplemental Table S2**). Prior reports have shown OPCs are dependent upon PDGF stimulation to maintain their proliferative capacity and prevent OL differentiation^42^, however the role of PDGFB in these processes remains less well studied. In our scATAC-seq datasets, we identified one OPC cluster (OPC-2) representing nearly 40% of all cells PDGFB^wt^ mice and less than 5% of PDGFB^ΔNLS^ cells (**Figs. 3c–d**; **Supplemental Tables S3-S4**), suggesting OPC-2 cells result from the expression of PDGFB and its capacity to localize to the nucleus. Conversely, two OL clusters (OL-1 and OL-2) were associated with of PDGFB^ΔNLS^ mice, showing minimal contributions from PDGFB^wt^ and non-tumor cells. These observations suggest that the OL-1 and OL-2 lineages are unique to the PDGFB^ΔNLS^ context, whereas OPC-2 cells are largely PDGFB^wt^-specific and implicate nuclear PDGFB as a regulator of OL differentiation from OPCs. Specifically, motif analysis and transcription start site (TSS) enrichment scoring revealed that chromatin from OPC-2 cells were enriched for centromere protein B (CENPB) binding sites while also demonstrating a largescale reduction in chromatin accessibility when compared to all other scATAC-seq cell type clusters (**Fig. 3e**; **Fig. S4d**). GO analysis of scRNA-seq differentially expressed genes (DEGs) revealed PDGFB^ΔNLS^-expressing cells upregulated gene sets pertaining to chromatin structure, centromeres and heterotopic nucleosome assembly including centromere protein A (CENPA) (**Fig. 3f**). These genomic and transcriptomic analyses suggest that nuclear PDGFB maintains OPC lineage by preventing OL differentiation and may do so by both restricting chromatin accessibility and altering centromeric heterochromatin.

**Figure 3.**
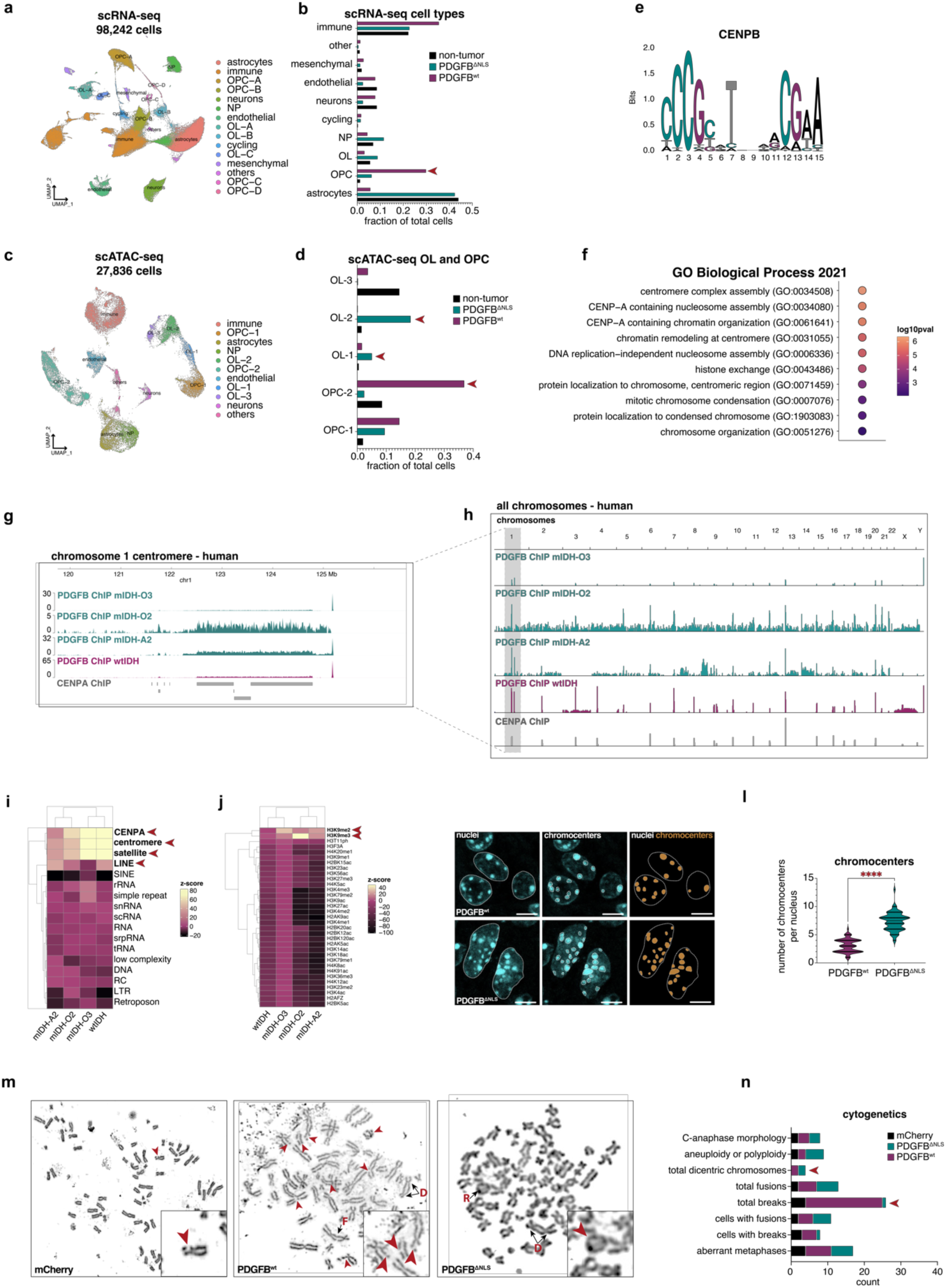
PDGFB binds heterochromatin at the centromere. (a) Dim plot of 98,242 cells from our scRNA-seq dataset comprised of PDGFB^wt^ (n=3), PDGFB^ΔNLS^ (n=3) and non-tumor (n=3) mouse brains showing cell type annotated clusters. (b) Bar plot of scRNA-seq cell types shows OPCs are enriched in PDGFB^wt^ tumors. (c) Dim plot of 27,836 cells from our scATAC-seq dataset comprised of PDGFB^wt^ (n=1), PDGFB^ΔNLS^ (n=1) and non-tumor (n=1) mouse brains showing cell type annotated clusters. (d) Bar plot of OPC and OL lineage clusters from scATAC-seq showing the OPC-2 lineage is predominantly found in PDGFB^wt^ tumors whereas OL-1 and OL-2 lineages are specifically enriched in PDGFB^ΔNLS^ tumors. (e) Motif plot from scATAC-seq showing CENPB binding sites are differentially accessible in PDGFB^wt^ OPC-2 cells. (f) Dot plot of GO analysis performed on scRNA-seq DEGs shows PDGFB^ΔNLS^-expressing cells are enriched for centromere-, nucleosome- and chromatin-related pathways when compared to PDGFB^wt^-expressing cells (DEGs: *p*<0.05, log2FC>0.05; GOs: *p*<0.05, log2FC>0.05). (g–h) Representative coverage plots of PDGFB ChIP-seq from human mtIDH (n=3) and wtIDH (n=1) glioma shows PDGFB peaks overlap with CENPA ChIP-seq peaks at centromeres. Peaks are shown for *(g)* the centromere of chromosome 1 and *(h)* select chromosomes. (i–j) Enrichment analysis of PDGFB ChIP-seq from human glioma samples shows wtIDH samples have the highest enrichment of PDGFB binding to *(i)* CENPA-binding sites, centromeres, satellite DNA regions, long interspersed nuclear elements (LINEs) and *(j)* H3K9me. (k) Representative confocal images of Hoechst-stained nuclei from PDGFB^wt^ and PDGFB^ΔNLS^ showing larger nuclei with more chromocenters in PDGFB^ΔNLS^ tumors; scale bar = 5 microns. (l) Violin plot showing increased chromocenter counts per cell in PDGFB^ΔNLS^. (m) Representative metaphase spreads of wtIDH GSCs (8-11) expressing mCherry, PDGFB^wt^ or PDGFB^ΔNLS^ lentivirus. Cytogenetic features of genome instability are highlighted: chromosome breaks (red arrows), fusions (F), rings (R) and dicentric chromosomes (D); n=35 metaphase spreads were analyzed per experimental cohort. (n) Quantification of cytogenetic genomic instability features from metaphase spreads shows that increased chromosome breaks are found in PDGFB^wt^ cells and that dicentric chromosomes are found in both PDGFB^wt^ and PDGFB^ΔNLS^ cells. H&E: hematoxylin and eosin; HGG: high-grade glioma; HR: hazard ratio; LGG: low-grade glioma. ChIP: chromatin immunoprecipitation; DEG: differentially expressed gene; GSC: glioma stem cell; GO: gene ontology; TSS: transcription start site; mtIDH-A2: IDH mutant astrocytoma grade II; mtIDH-O2: mutant oligodendroglioma grade II; mtIDH-O3: mutant oligodendroglioma grade III; wtIDH: IDH wild-type.

### Reduced binding of PDGFB to centromeric heterochromatin leads to genome instability

The forgoing data suggest an association exists between nuclear PDGFB and centromeres. To examine the nature of this putative relationship, we performed PDGFB ChIP-sequencing (ChIP-seq) on mouse and human glioma samples. Across all human and mouse samples analyzed, the strongest PDGFB ChIP-seq signals were found at the centromere, which included CENPA sites, long interspersed nuclear elements (LINEs), and repeat-rich satellite DNA regions marked by repressive histone 3 lysine 9 methylation (H3K9me)^43–45^ (**Figs. 3g–j**; **Figs. S4e–f**). In human glioma samples, PDGFB ChIP-seq enrichment at these loci was lowest in a recurrent mtIDH tumor and was highest in a wtIDH tumor, which suggests that loss of centromeric binding in mIDH tumors may contribute to disease progression. To independently validate our observation that PDGFB binds the centromere, we employed high resolution microscopy of PDGFB^wt^ or PDGFB^ΔNLS^ nuclei to quantify chromocenters, regions of densely-packed centromeric and pericentromeric heterochromatin^46–49^ that are readily visible in murine nuclei. We observed that PDGFB^wt^ nuclei contained less chromocenters than PDGFB^ΔNLS^ nuclei, suggesting that aberrant levels of PDGFB in the nucleus influence centromeric heterochromatin (**Figs. 3k–l**). These analyses also revealed that PDGFB^wt^ nuclei were smaller and rounder than PDGFB^ΔNLS^ nuclei, the latter of which presented with atypical morphologies (**Fig. S4g**).

These interactions with centromeres, coupled with our mass spectrometry data showing that PDGFB associates with histones, led us to examine whether PDGFB alters histone-DNA interactions by binding directly to nucleosomes. To assess this, we performed a nucleosome extraction assay and found that PDGFB was present in mononucleosome preparations (**Figs. S4h–i**) and was associated with a reduction in histone H3 in PDGFB^ΔNLS^ nucleosomes (**Figs. S4j– k**). Consistent with these data and our single cell analyses, we found that *CENPA* expression was higher in PDGFB^ΔNLS^ mice and conversely, that PDGFB^wt^ mice showed enrichment for *CENPB* expression (**Fig. S4l**). These results show that nuclear localization of PDGFB is important for determining centromeric heterochromatin states. One consequence of aberrant centromere formation is genomic instability resulting from chromosome missegregation, aneuploidy, chromosomal rearrangements and micronucleus formation^34,50,51^. Accordingly, we next examined whether PDGFB^wt^ and PDGFB^ΔNLS^ contributed to genomic instability in mouse tumors and human glioma stem cell lines. We inferred CNVs using our mouse scRNA-seq dataset and observed frequent chromosome 1 amplifications in PDGFB^wt^ tumor cells (**Fig. S4m**) In human GSCs, we used mitotic spreads to assess chromosomal integrity, finding that PDGFB^wt^ led to increased chromosome breaks whereas PDGFB^ΔNLS^ resulted in more frequent chromosomal fusion events (**Figs. 3m–n**). Both PDGFB^wt^ and PDGFB^ΔNLS^ led to the presence of dicentric chromosomes having two centromeres that were absent from controls, demonstrating that appropriate levels of nuclear PDGFB are required for proper positioning of the centromere. Collectively, these studies suggest that elevated nuclear PDGFB is a driver of chromosomal breaks and suggest that insufficient nuclear PDGFB may increase chromosomal fusions. Moreover, these experiments demonstrate that PDGFB is important for chromatin architecture and genomic stability, which is likely accomplished via direct interactions between PDGFB and nucleosomes.

### Mutant IDH impairs PDGFB chromatin binding

Given that high PDGFB expression is associated with poor survival outcomes in mIDH glioma, we sought to define how mutations to IDH alter PDGFB-chromatin interactions and to discern the effects of these alterations on gliomagenesis and progression. To do this, we overexpressed PDGFB and IDH1^R132H^ simultaneously in our piggyBac IUE model (PDGFB^wt^–IDH1^R132H^), which conferred reduced survival times when compared to PDGFB^wt^ overexpression alone (**Figs. 4a– b**). Unexpectedly, PDGFB^wt^–IDH1^wt^ co-overexpression trended towards worse survival outcomes than PDGFB^wt^–IDH1^R132H^ mice, whereas neither IDH1^R132H^ nor IDH1^wt^ overexpression alone were able to induce tumorigenesis in the absence of PDGFB expression. These data suggest that high protein levels of IDH1 are oncogenic in the context of PDGFB and confirm that gliomagenesis in mice requires an additional event secondary to the IDH1^R132H^ mutation. Indeed, total IDH1 (IDH1^wt^ and IDH1^R132H^) protein levels were higher in mIDH cell lines when compared to wtIDH cells (**Fig. 4c**) and *IDH1* and *IDH2* expression was highest in PDGFRA+ tumor cells, which are the majority of glioma cells in mIDH tumors and a small percentage of GSCs in wtIDH tumors (**Fig. 4d**; **Fig. S5a**). Transcriptomic data from the Allen Brain Atlas confirmed PDGFRA+ OPCs are enriched for *IDH1* and *IDH2* expression (**Fig. 4e**; **Fig. S5b**), suggesting that OPCs may be particularly susceptible to the effects of IDH somatic mosaicism. To investigate the effects of IDH1^R132H^ on PDGFB localization, we overexpressed PDGFB^wt^ in mIDH GSC cell lines derived from mIDH patients and compared cell fractions via immunoblotting. In contrast to wtIDH GSCs where overexpression of PDGFB^wt^ led to an enrichment of PDGFB in the chromatin fraction, overexpression in mtIDH GSCs, PDGFB^wt^ was largely absent from the chromatin fraction, similar to the effects of PDGFB^ΔNLS^ overexpression (**Fig. 4f**). Immunostaining of PDGFB^wt^–IDH1^R132H^ and PDGFB^wt^–IDH1^wt^ mice confirmed nuclear PDGFB was reduced in tumor cells when IDH1^R132H^ was co-overexpressed with PDGFB^wt^ whereas PDGFB was increased when IDH1^wt^ was present; both IDH1^wt^ and IDH1^R132H^ variants could be found in the nucleus (**Fig. 4g**; **Fig. S5c**), which is consistent with prior reports documenting non-classical roles for biochemical enzymes in the nucleus^52^. Moreover, in human mIDH tumors we found that CENPB expression was upregulated in IDH1^R132H^– tumor cells when compared to IDH1^R132H^+ cells (**Fig. S5d**). These observations suggest that the presence of IDH1^R132H^ impairs CENPB expression, which may destabilize the centromere. Our collective data suggest that high levels of IDH1^wt^ increase PDGFB nuclear localization even more than PDGFB overexpression alone, thereby accelerating PDGFB^wt^ phenotypes caused by elevated nuclear PDGFB. In mIDH glioma, high levels of IDH1^R132H^ decrease nuclear localization of PDGFB, rendering PDGFB unable to bind centromeric heterochromatin, perhaps through changes to CENPB binding. Taken together, our experimental evidence suggests IDH1^R132H^ mutations mechanistically serve to destabilize the centromere by impeding PDGFB-chromatin interactions, which in turn may facilitate gliomagenesis. These data propose a new mechanism for IDH mutations in glioma as inhibitors of PDGFB binding to chromatin.

**Figure 4.**
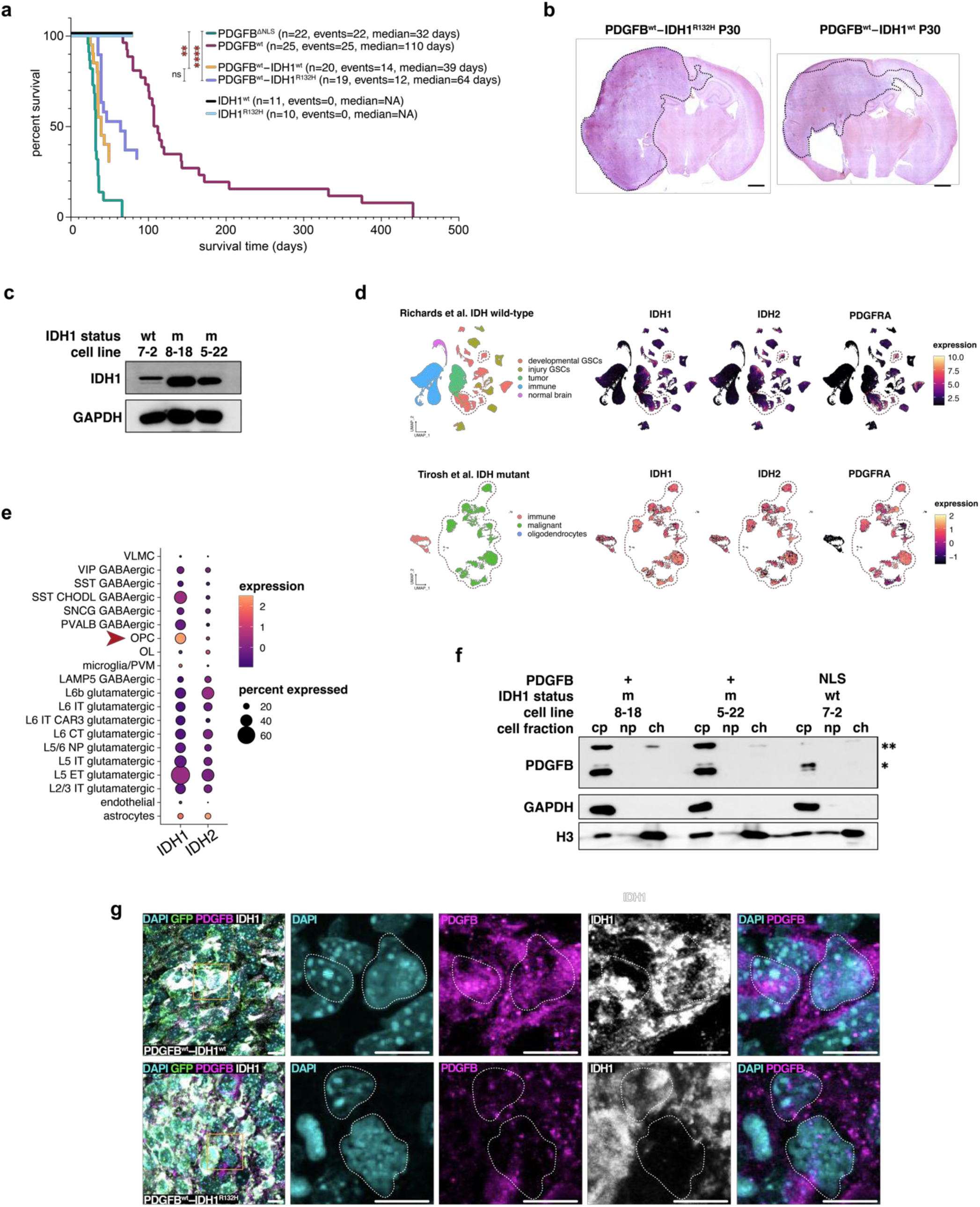
IDH1R132H impairs nuclear translocation of PDGFB. (a) Kaplan-Meier survival analysis of PDGFB^wt^ (n=25, median=110 days), PDGFB^ΔNLS^ (n=22; median=32 days), IDH1^R132H^ (n=10; median=NA), IDH1^wt^ (n=11; median=NA), PDGFB^wt^–IDH1^R132H^ (n=19; median=64 days) and PDGFB^wt^–IDH1^wt^ (n=20; median=39 days) mice; pairwise *p*-values are denoted in the figure. (b) H&E staining of a coronal section of end-stage PDGFB^wt^–IDH1^wt^ (P30) and PDGFB^wt^– IDH1^R132H^ (P30) tumors. Black dashed line denotes tumor; scale bar = 2 mm. (c) Immunoblot of total IDH1 protein in wtIDH (7-2) and mtIDH (8-18; 5-22) GSCs shows mIDH cell lines have increased total IDH1 protein. (d) Dim plot and corresponding feature plots are shown for scRNA-seq datasets published by Richards et al. and Tirosh et al. for human glioma. Feature plots show *IDH1* and *IDH2* are highly expressed in mIDH tumor cells, which also express *PDGFRA* (red dashed lines). In wtIDH glioma, PDGFRA+ tumor cells and PDGFRA+ GSCs (red dashed lines) show high expression of *IDH1* and *IDH2*. (e) Dot plot of human brain scRNA-seq data from Allen Brain Atlas shows *IDH1* expression is highest in OPCs. (f) Immunoblots of cell fractions from primary GSCs derived from two IDH1^R132H^+ mIDH glioma patient samples (8-18; 5-22). PDGFB translocation to the nucleus is reduced when PDGFB^wt^ is overexpressed in mIDH GSCs, which is phenotypically similar to overexpression of PDGFB^ΔNLS^ in wtIDH GSCs (7-2); * = 54.6kD; ** = 18.4kD; ch: chromatin fraction; np: nucleoplasm; cy: cytoplasmic fraction. (g) Immunostaining of PDGFB^wt^–IDH1^wt^ and PDGFB^wt^–IDH1^R132H^ mouse tumors shows reduced nuclear PDGFB in tumor cells (white dashed lines) in PDGFB^wt^–IDH1^R132H^ mice as compared to PDGFB^wt^–IDH1^wt^ mice; scale bar = 10 microns.

### PDGFB is immune–derived in human glioma

While examining human glioma scRNA-seq datasets^41,53,54^, we found that expression of PDGFB was largely absent from tumor cells, which were frequently PDGFRA+ in mIDH tumors (**Figs. 5a– c**). Instead, we observed that in both tumor and non-tumor contexts, PDGFB was most highly expressed by immune cells and was mainly present in myeloid-derived CD11b+ macrophages. Immunostaining of IDH1^R132H^ and PDGFB in mIDH glioma samples confirmed that cytoplasmic staining of PDGFB was absent from tumor cells but was detected in CD11b+ myeloid (**Figs. 5d– e**). These observations suggested to us that immune-derived PDGFB is localizing to the nuclei of OPCs to promote OPC proliferation and inhibit differentiation of tumor cells. To test this, we overexpressed PDGFB *in vitro* in primary mouse microglia and cultured wtIDH GSCs in the microglia-conditioned media (**Fig. 5f**). Immunoblotting of GSC cell fractions confirmed that PDGFB localized to the chromatin fraction, demonstrating that exogenous PDGFB can localize to the nucleus and bind chromatin in glioma cells bearing the cognate receptor, PDGFRA (**Fig. 5g**). These experimental results implicate immune cells as the origin of PDGFB in human glioma and demonstrate that a mammalian protein produced by one cell can translocate to the nucleus of a different cell.

**Figure 5.**
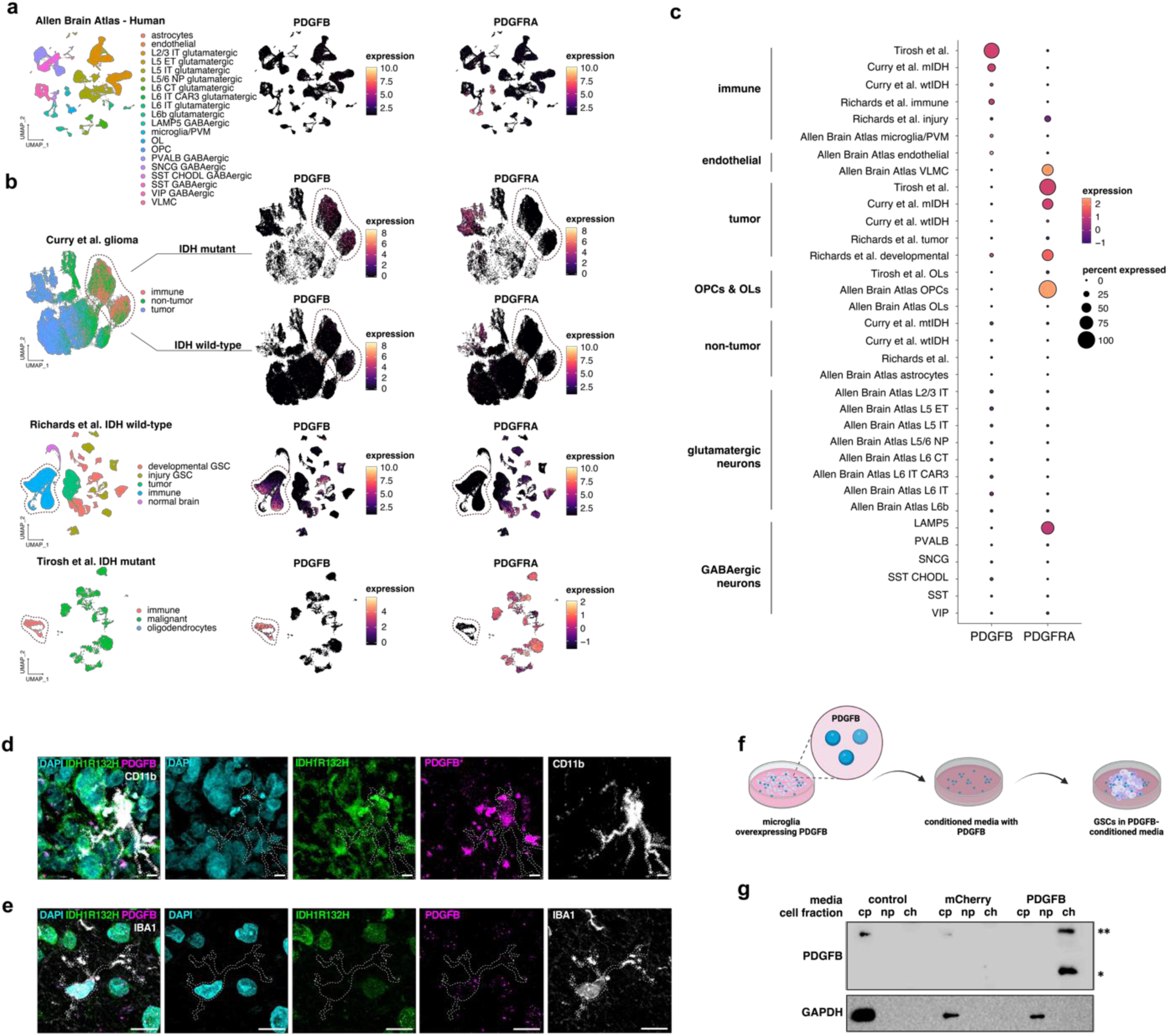
PDGFB is produced by immune cells in glioma. (a–b) Dim plots and corresponding feature plots for human scRNA-seq datasets from normal adult cortex *(a)* and glioma *(b)* showing *PDGFB* expression is detected in microglia and macrophages whereas *PDGFRA* is detected in OPCs and glioma cells. (c) Dot plot of human scRNA-seq datasets in *(a)* and *(b)* showing comparative levels of *PDGFB* and *PDGFRA* across non-tumor and glioma datasets. Note that *PDGFB* and *PDGFRA* expression occur in mutually-exclusive cell types. (d–e) Immunostaining of surgically-resected human mIDH glioma sample shows PDGFB is minimally expressed in IBA1+ microglia *(d)* and is highly expressed in CD11b+ myeloid cells *(e)*; scale bar = 10 microns. (f) Schematic showing experimental setup of PDGFB conditioned media experiment. (g) Immunoblots of cell fractions from wtIDH GSCs (7-2) cultured in microglia-conditioned media shows PDGFB is detected in the nucleus of GSCs cultured in PDGFB^wt^-overexpressing microglia.

## Discussion

Our studies identified high levels of PDGFB in human mIDH glioma and found PDGFB binds chromatin in human tumors. We developed a novel model of immunocompetent glioma, and found find that PDGFB normally translocates to the nucleus to bind the centromere. We propose that decreased levels of nuclear PDGFB engenders genome instability resulting from improper centromere formation and maintenance. Our experiments revealed that mutant IDH restricts nuclear localization of PDGFB and consequently, centromere binding by PDGFB is reduced. Our work shows that PDGFB and mutant IDH synergize to accelerate gliomagenesis in the absence of other mutations, suggesting that mutant IDH and high levels of PDGFB are sufficient to induce glioma. Finally, we identified the immune compartment as the predominant source of PDGFB in mIDH glioma, confirming that immune-derived PDGFB can translocate to the nucleus of glioma cells and bind chromatin. These studies identify new roles for growth factors in the nucleus as important chromatin remodelers and advance our understranding of how mutations to IDH can engender glioma.

### Mutant IDH impairs chromosome stability

In contrast to wtIDH glioma, mIDH tumors occur in patients less than 40 years of age^55–58^ and frequently lack mutations to tumor suppressor genes or oncogenes that drive wtIDH glioma. Accordingly, the role of IDH mutations in tumor initiation is controversial and conclusive evidence describing how they autonomously contribute to human gliomagenesis remains obscure. Alongside prior reports demonstrating IDH mutations in glia are detected in the healthy human brain^59^ and that IDH1^R132H^ is insufficient to drive gliomagenesis alone, our experimental findings suggest mIDH may serve as a predisposition to cancer rather than a *bona fide* driver event. Our data support a model in which IDH1^R132H^ impairs PDGFB binding to heterochromatin in PDGFRA-expressing cells, which includes OPCs – the most proliferative cells in the adult human brain^60–62^ (**Fig. 6a**). When considered collectively, our experimental findings suggest that in proliferating OPCs, PDGFB must enter the nucleus and bind centromeric heterochromatin to ensure faithful segregation of the genome. Moreover, our work suggests that IDH levels may impact the localization of PDGFB, although direct interactions between IDH and PDGFB were not observed from our mass spectrometry or CoIP experiments (data not shown). OPCs, which show a unique enrichment in IDH expression, may be particularly susceptible to the effects of IDH mutations, the effect of which greatly reduces translocation of PDGFB to the nucleus. We postulate that when excess nuclear PDGFB is present, chromosomes are susceptible to breakage due to ectopic centromere formation caused by too much PDGFB. Conversely, when insufficient nuclear PDGFB is present, improper centromere formation may lack the required constituents to segregate properly. These data have led us to propose a two-step model of mIDH gliomagenesis that requires both (1) mutant IDH and (2) a source of elevated PDGFB (**Fig. 6b**). In this model, IDH mutations prevent PDGFB from localizing to the nucleus, leading to chromosome destabilization and aneuploidy. Given our data showing PDGFB is sufficient to induce genomic instability in the absence of mutations, future studies should aim to discern whether aberrant PDGFB-chromatin interactions can serve as an initiating event to induce canonical IDH mutations in the absence of other genomic drivers.

**Figure 6.**
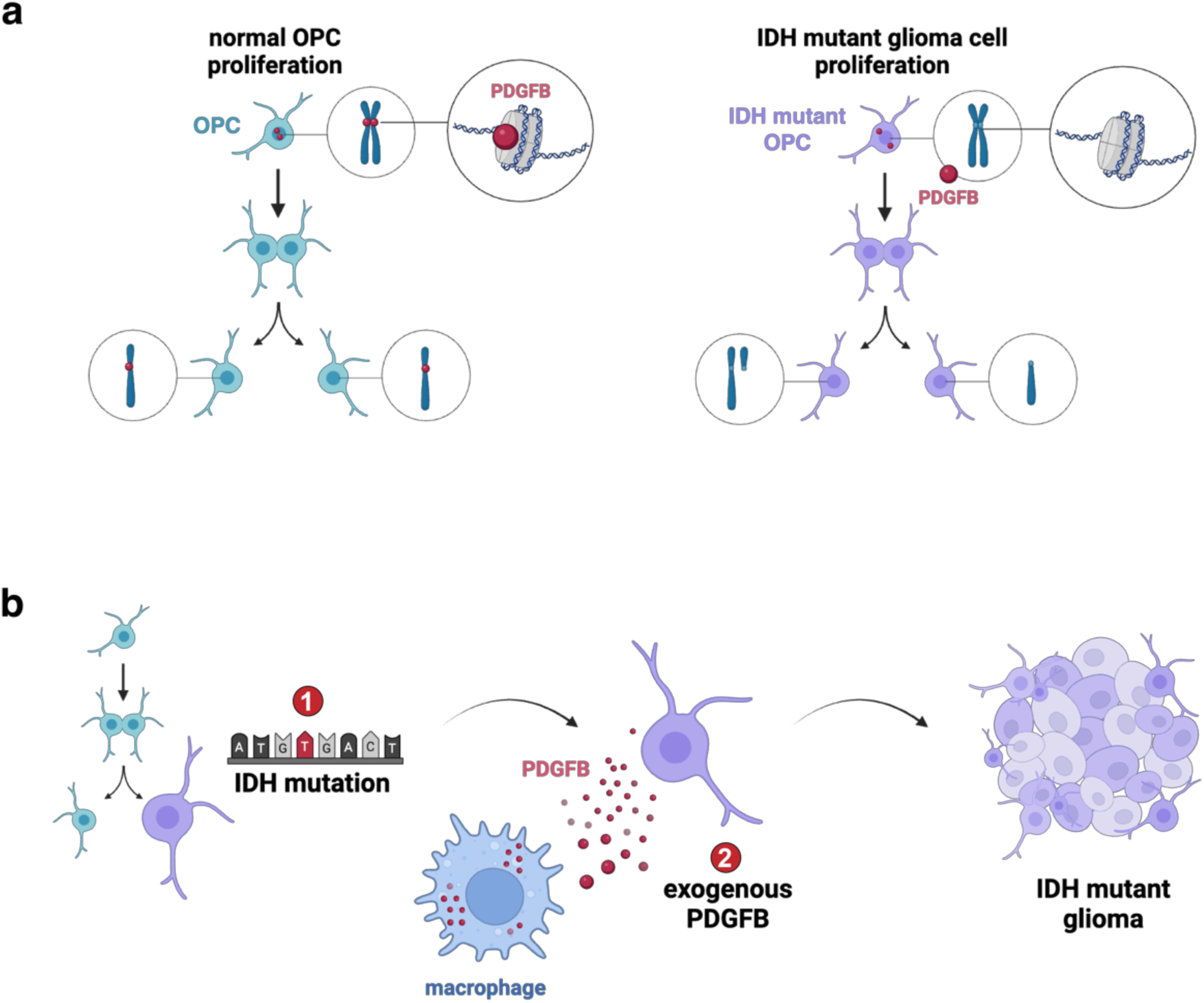
Working model of nuclear PDGFB in glioma. (a) Schematic showing a working model of nuclear PDGFB. In non-tumor OPCs, PDGFB binds centromeric DNA to facilitate proper chromosome segregation. In mIDH glioma, immune-derived PDGFB is unable to bind the centromere, leading to genomic instability. (b) A two-step model for mIDH gliomagenesis: (1) the IDH1 mutation, and (2) a source of constitutive PDGFB signaling.

### The neuroimmune niche is the source of PDGFB in mIDH glioma

Our research identified immune cells as a primary source of PDGFB in both humans and mice. In the non-tumor brain, PDGFB is mainly produced by microglia and perivascular macrophages, which represent the myeloid compartment of the central nervous system^63^ and is also produced in lesser quantities by vascular cells. In glioma, increased production of PDGFB is attributed to expansion of the myeloid compartment, which is populated by tumor-infiltrating myeloid cells. The observations that PDGFB is largely immune-derived, sufficient to generate glioma in the absence of mutations^64,65^ and synergizes with IDH1^R132H^ to accelerate disease progression raise the possibility that exogenous PDGFB may contribute to gliomagenesis. Indeed, chronic inflammation has been linked to an increased risk of certain cancers, including esophageal, lung and colorectal cancer^66–68^, suggesting a causative role for inflammatory processes in human oncogenesis. In addition to inflammation, viral infections have been established as causative drivers of cancer, although evidence for this in the context of neurological neoplasms is lacking^69–82^. Despite an absence of conclusive studies linking viral infections to brain tumors, an abundance of correlative evidence suggests that a viral origin for certain brain tumors may exist^83–86^, including the identification of the PDGFB homolog v-*sis*, which drives glioma in marmosets and shares 92% homology with PDGFB^87–92^. While future research endeavors will need to examine whether inflammatory or viral inputs engender gliomagenesis in humans by providing a constitutive source of PDGFB, our data implicate microenvironmental constituents as potentially important contributors to the initiation of brain tumors.

### Paracrine-mediated remodeling of chromatin states in the human brain

An unexpected finding of this research study was the discovery that GSCs can internalize exogenous PDGFB and translocate it to the nucleus. Nearly 40 years ago, researchers demonstrated that growth factors, including PDGFB, translocate to the nucleus to bind chromatin in cells bearing the appropriate surface receptors, like PDGFRA^16^. Several other studies reported similar findings for epidermal growth factor, nerve growth factor, gonadotropin and angiotensin II, however, these findings were controversial given the conceptual framework mandating lysosomal degradation of all endocytosed proteins^93^. Indeed, the implications of paracrine-mediated chromatin remodeling, wherein chromatin is directly bound and regulated by proteins produced exogenously, could be far-reaching. Our current understanding of receptor-ligand interactions, including receptor tyrosine kinases like PDGFRA, postulates that transcriptional changes to target cells are mediated entirely through phosphorylation-dependent cascades that are triggered by binding of a ligand to the receptor at the cell surface. This binding activates-complex series of events that culminates in the translocation of endogenous cytoplasmic proteins to the nucleus. Undoubtedly, membrane-bound receptor-ligand interactions are potent mediators of paracrine-mediated cell differentiation, proliferation and growth that have documented roles in both developmental and oncogenic processes. Our work builds upon these principles of molecular biology to suggest that paracrine signaling paradigms may also exist in which production of proteins from one cell can be internalized by another cell and subsequently undergo nuclear localization. Given the abundance of receptor-ligand interactions in myriad biological processes, future research should aim to better define whether additional functions exist for these proteins in the nucleus.

## Acknowledgements

This work was supported by grants from the NIH (R35-NS132230 to B.D., R01NS124093 to B.D. and G.R., R01CA223388 to B.D., U01CA281902 to B.D., R01NS094615 to G.R.). R.N.C. is supported by grants from the NIH (F31CA265156 and F99CA274700). This study includes work performed at: (1) the Single Cell Genomics Core at BCM partially supported by NIH grant S10OD025240 and CPRIT RP200504; and (2) the Cytogenetics and Cell Authentication Core at MD Anderson Cancer Center.

## Contributions

R.N.C., B.D., and G.R. are responsible for the conception of this project, study design and interpretation of results. R.N.C. prepared all IP-MS, scRNA-seq and scATAC-seq samples from both human and mouse. P.H. performed all Western blotting. R.N.C., M.F.M and P.H. performed all mouse surgeries. R.N.C., M.F.M., Y.K., J.M., L.F., N.P. and I.O. performed immunostaining, imaging and imaging analyses; J.L. provided microscopy equipment and resources. B.L. and D.S. performed all ChIP and ChIP-seq experiments. U.K. and P.H. performed cloning and generated lentivirus. D.J.C. and P.H. performed all *in vitro* experiments. J.M. and A.M. performed cytogenetics analyses. R.N.C., S.W., M.G., A.O.H. and A.S.H. performed bioinformatics analyses on human and mouse scRNA-seq, scATAC-seq and ChIP-seq datasets. P.H. and M.F.M. maintained all mouse colonies and assisted with tissue harvesting and fixation. C.M. performed histopathological diagnoses of mouse tumors. R.N.C., B.D. and G.R. prepared the manuscript with feedback from all authors.

## RESOURCE AVAILABILITY

### Lead Contact

Further information and requests for resources and reagents should be directed to and will be fulfilled by the Lead Contact, Ganesh Rao.

### Materials Availability

All published reagents will be shared on an unrestricted basis after completion of a material transfer agreement; reagent requests should be directed to the lead contact.

## Data and Code Availability

The RNA-seq dataset generated during this study is available at the NCBI GEO website.

### Experimental Methods

#### Human data

Adult patients at St. Luke’s Medical Center and Ben Taub General Hospital provided preoperative informed consent to participate in the study and gave consent under Institutional Review Board Protocol H35355. Patients included males and females. Clinical characteristics were maintained in a deidentified patient database and are summarized in **Table S1**.

Tumor samples were collected during surgery and immediately placed on ice. Tissue was divided for use in subsequent genomic, histopathological, proteomic or biochemical studies. Patient samples were collected separately for pathology and molecular subtyping. Histopathology and molecular subtyping of IDH and chromosome 1p/19q deletion status were confirmed by board-certified pathologists.

#### Histology

Human samples were retrieved from the operating room on ice and then fixed in 4% paraformaldehyde in PBS for 12 hr at 4°C before being transferred to 70% EtOH. Mouse samples were retrieved following euthanasia and fixed according to the same protocol. Paraffin embedding was performed by the Breast Cancer Pathology Core at Baylor College of Medicine. All human and mouse specimens were evaluated by a board-certified neuropathologist according to current guidelines and standard practices.

#### Immunostaining

For immunostaining, 10−μm paraffin-embedded human glioma sections were cut, deparaffinized and subject to heat-induced epitope retrieval (HIER) using antigen retrieval buffer (10 mM sodium citrate, 0.05% Tween 20, pH 6.0) when needed. Sections were blocked for 1 hr at room temperature and kept in primary antibody incubation overnight at 4°C.

Species-specific secondary antibodies tagged with Alexa Fluor corresponding to emission spectra 488 nm, 568 nm, or 647 nm (1:1,000, Thermofisher) were used for immunofluorescence and Hoechst nuclear counter staining (1:50,000; Thermofisher, H3570) was performed before coverslipping with Vectashield antifade mounting medium (Vector Laboratories, H-1000).

#### piggyBac *in utero* electroporation

Tumor mice were generated by single-sided intraventricular injection of overexpression constructs targeting Glast-expressing mouse neural precursor cells (NPCs) via piggyBac transposase technology. Surgery and *in utero* electroporation were performed on CD1 wildtype pregnant damns at E16.5 according to our previously established protocols^94^. PDGFB^wt^, PDGFB^ΔNLS^, IDH1^wt^, IDH1^R132H^, PDGFB^wt^-IDH1^wt^ and PDGFB^wt^-IDH1^R132H^ mice were generated using piggyBac constructs driving overexpression of the full PDGFB CDS or PDGFB with modifications to the C-terminus at amino acids 214-227: ^213^RVRRPPKGKHRKFKHTH^231^ converted to ^213^SVSSPPSGSHSSFSHTH^231^. All mice received co-electroporation of a piggyBac construct driving overexpression of GFP. Tumor brains were collected from mice either at matched time points or end-stage disease. Mice were monitored for symptoms indicative of tumor burden, including lethargy, hunched posture, decreased appetite, poor grooming maintenance, squinting of the eyes, partial limb paralysis and/or ataxia, denoting the IACUC permitted endpoint.

#### Immunoprecipitation mass spectrometry (IP-MS)

Microdissected GFP-expressing mouse tissue were washed with cold PBS and dissociated using a pellet homogenizer. Cell lysates were prepared in coimmunoprecipitation (CoIP) buffer (50 mM Tris-HCl, 120 mM NaCl, 1 mM EDTA, 0.5% NP-40) through ultracentrifugation 100,000 g for 30 min at 4°C. Supernatant was harvested and incubated with Pierce Protein A/G Magnetic Beads (Thermofisher, 88802) for 30 min at 4°C. The precleared lysate was isolated and incubated with anti-mouse IgG (Santa Cruz, sc-2025) for 30 min at 4°C before being incubated again with Pierce Protein A/G Magnetic Beads for another for 30 min at 4°C. The precleared, IgG-immunoprecipitated lysate was then incubated with mouse anti-PDGFB (F-3) antibody (Santa Cruz, sc-365805) overnight at 4°C. All beads were collected using a magnetic separator, washed briefly in CoIP buffer, boiled in 2× NUPAGE LDS sample buffer (Life Technologies), and resolved on a NuPAGE 10% Bis-Tris Gel (Life Technologies). The eluted proteins were visualized with Coomassie Brilliant blue stain, excised into gel pieces according to the molecular weight, and in-gel digested with trypsin. The liquid chromatography-tandem mass spectrometry (MS) analysis was carried out using the nanoLC1000 system coupled to an Orbitrap Fusion mass spectrometer (ThermoScientific). The peptides were loaded on a two-column setup with a precolumn (2 cm × 100 µm internal diameter [I.D.]) and analytical column (5 cm × 150 µm I.D.); filled with Reprosil-Pur Basic C18 (1.9 µm; Dr. Maisch GmbH). The peptide elution was done using a discontinuous gradient of 90% acetonitrile buffer (B) in 0.1% formic acid (5 to 28% B, 800 nL/min: 45-min gradient). The MS instrument was operated in data-dependent mode with MS1 acquisition in Orbitrap (120,000 resolution, AGC 5e5, 50-ms injection time) followed by MS2 in Ion Trap (Rapid Scan, HCD 30%, AGC 5e4). The MS raw data were searched using Proteome Discoverer 2.0 software (ThermoScientific) with Mascot algorithm against the mouse National Center for Biotechnology Information (NCBI) refseq database updated 24 March 2020. The precursor ion tolerance and product ion tolerance were set to 20 ppm and 0.5 Da, respectively. A maximum cleavage of two with trypsin enzyme, dynamic modification of oxidation on methionine, protein N-term acetylation, and destreak on cysteine were allowed. The peptides identified from the mascot result file were validated with FDR<0.05. The gene product inference and quantification were done with the label-free iBAQ approach using the gpGrouper algorithm. For statistical assessment, missing value imputation was employed through sampling a normal distribution N (μ-1.8 σ, 0.8σ), where μ, σ are the mean and standard deviation of the quantified values. For differential analysis, we used the moderated t test as implemented in the R package limma, and multiple-hypothesis testing correction was performed with the Benjamini-Hochberg procedure. All proteins with a log2(fold change)>2 and p<0.05 were considered for analysis.

#### Western blotting

Human and mouse lysates were prepared as described above using CoIP or RIPA buffer. Proteins were analyzed by Western blotting using 12% SDS polyacrylamide gels, followed by wet transfer to nitrocellulose membrane at 400 mA for 1 hr. Membranes were blocked by EveryBlot Blocking Buffer (BioRad, 12010020) or 5% milk diluted in Tris-buffered saline supplemented with Tween20 (TBST) followed by overnight incubation at 4°C in primary antibody. Membranes were washed TBST and incubated at room temperature for 1 hr in horseradish peroxidase-conjugated secondary antibodies, washed again with TBST, and developed using luminol reagent.

#### Cell fraction

Glioma stem cell spheres were collected and washed with PBS, resuspended with 100μl cytoplasmic lysis buffer (0.15% (vol/vol) NP-40, 10 mM Tris-HCl (pH 7.0), 150 mM NaCl) supplemented with 1× protease inhibitor mix and incubated on ice for 10 min. Cell lysate were layered onto 500μl sucrose buffer (10 mM Tris-HCl (pH 7.0), 150 mM NaCl, 25% (wt/vol) sucrose) supplemented with 1× protease inhibitor mix and centrifuged for 10 min at 16,000 *g* at 4°C. The top liquid layer was transferred to a new tube and stored on ice as the cytoplasmic fraction. The nuclei pellet was washed with 800μl nuclei wash buffer (0.1% (vol/vol) Triton X-100, 1mM EDTA) supplemented with 1× protease inhibitor mix. Nuclei were resuspended with 50μl glycerol buffer (20 mM Tris-HCl (pH 8.0), 75 mM NaCl, 0.5 mM EDTA, 50% (vol/vol) glycerol, 0.85 mM DTT) supplemented with 1× protease inhibitor mix and mixed with 50μl nuclei lysis buffer (1% (vol/vol) NP-40, 20 mM HEPES (pH 7.5), 300 mM NaCl, 1M urea, 0.2 mM EDTA, 1mM DTT) supplemented with 1× protease inhibitor mix and incubated on ice for 10 min after mixing well. Samples were spun at 18,500 *g* for 10 min at 4°C. Supernatant was collected as the nucleoplasm fraction and the DNA pellet was collected as the chromatin fraction. All cell fractions were stored at −80°C until further use.

For nucleosome extractions, cells were maintained in culture, harvested and pelleted. Nucleosomes were prepared using the Nucleosome Preparation Kit (Active Motif, 53504) per the manufacturer’s instructions.

#### Primary microglia culture

Primary microglia were obtained from mixed glia cultures grown from forebrains of P1 ICR mouse pups. Briefly, forebrains were isolated and homogenized into single-cell suspensions by triturating with fire-polished Pasteur pipettes in Dulbecco’s modified Eagle’s medium (DMEM) with high glucose and HEPES (GenDEPOT, CM003-050) containing penicillin/streptomycin (Gibco, 15080-063), GlutaMAX (Gibco, 35050061), 10% heat-inactivated fetal bovine serum (FBS; GenDEPOT, F0601-050). Cells were plated in 75 cm^2^ T-flasks and incubated for 2 weeks. Floating microglia were detached from flasks by mild shaking, harvest and filtered through a 70-µm cell strainer to remove cell debris. Microglia were then plated in new 75 cm^2^ T-flasks for use in subsequent experiments.

#### Microglia-conditioned media

Lentivirus was added to primary microglia and cultured for 5 days until mCherry expression was observed. Primary culture media was removed, and microglia cells were washed with PBS. Neurobasal Media was added to microglia, incubated for 5 days, and collected as microglia-conditioned media. Microglia were collected and kept at −80°C for further analysis. 1×10^6^ glioma stem cells were added to conditioned media and cultured for 3 days. All glioma stem cells were collected and kept at −80°C for further analysis.

#### Lentivirus

HEK293T cells were cultured in DMEM + 10% FBS and transfected with PSPAX2, PMD2G, PHAGE-PDGFB^wt^, PHAGE-PDGFB^ΔNLS^, PHAGE-IDH1^wt^, PHAGE-IDH1^R132H^ or PHAGE-mCherry constructs using Lipofectamine3000 transfection reagent (Invitrogen, L3000-015). Conditioned media from transfected HEK293T cells were collected and kept at 4C. lentivirus were precipitated by PEG-itTM (SBI, LV825A-1) and redissolved in PBS. Lentivirus was aliquoted and kept at −80°C until time of use.

Primary glioma stem cell lines were cultured in Neurobasal Media [DMEM-F12 Thermofisher, 11320082) supplemented with B27 (×1; Thermofisher, no. 17504001), hBFGF (20 ng/ml; Sigma Aldrich, F0291) and EGF (20 ng/ml, Peprotech, no. 100-47)]. Cells were infected with lentivirus. Following 48 h of infection, cells were selected by puromycin (1 μg/ml; Thermofisher, A1113803).

#### Chromatin immunoprecipitation (ChIP)

Human and mouse tumor samples were collected according to the aforementioned protocols and were dissociated in cold PBS using a pellet homogenizer on ice. Chromatin was crosslinked using 1.1% formaldehyde solution at room temperature for 10 min, followed by addition of 0.1 M glycine. Cell pellets were collected by centrifugation at 3500 rpm for 5 min at 4°C, washed with PBS, and frozen at −80 °C until further processing. Pellets were resuspended with PBS/PMSF supplemented with 0.5% Igepal, washed with cold ChIP buffer (0.25% TritonX, 10 mM EDTA, 0.5 mM EGTA, 10 mM Hepes [pH 6.5]), lysed with ChIP lysis buffer (0.5% SDS, 5 mM EDTA, 25 mM Tris⋅HCl [pH 8]) for 20 min at room temperature. Lysates were sonicated to 250 to 350 bp using the Diagenode Bioruptor and chromatin shearing was validated using a TapeStation. Chromatin was quantified using the Quant-iT double-stranded DNA (dsDNA) assay kit (Thermofisher, Q33120), diluted five-fold (2 mM EDTA, 150 mM NaCl, 1% Triton X-100, 20 mM Tris⋅HCl; pH 8.0) and incubated with antibody overnight at 4°C with rotation. For ChIP-PDGFB, chromatin was first incubated with anti-mouse IgG (Santa Cruz, sc-2025), pulldown performed and then subsequently incubated with mouse anti-PDGFB (F-3) antibody (Santa Cruz, sc-365805) overnight at 4°C. Lysates were incubated with Protein A/G magnetic beads (Thermofisher, 88802) for 6 h at 4°C, followed by washing with Tris-SDS-EDTA-I buffer (0.1% SDS, 1% Triton X-100, 2 mM EDTA, 150 mM NaCl, 20 mM Tris⋅HCl; pH 8.0), Tris-SDS-EDTA-II buffer (TSEI buffer with 500 mM NaCl), LiCl buffer (250 mM LiCl, 1% Nonidet P-40, 1% deoxycholate, 1 mM EDTA, 10 mM Tris⋅HCl; pH 8.0), and Tris-EDTA buffer (10 mM Tris-HCl, 1 mM EDTA; pH 8.0). To release DNA fragments, samples were incubated in freshly prepared elution buffer (1% SDS, 0.1 M NaHCO3) for 20 min at 65°C twice. Elutions were treated with proteinase K (0.4 mg/mL; Thermofisher, AM2546) and NaCl (0.125M) overnight at 65°C for reverse crosslinking. Subsequently, ChIP-DNA was purified using a PCR purification kit (Qiagen, 28104) and quantified using the Quant-iT dsDNA assay kit. For ChIP-Seq experiments, 10 ng of ChIP-DNA was used for library preparation as described below.

#### ChIP-sequencing

ChIP libraries were prepared using the TruSeq ChIP Library Preparation Kit (Illumina, IP-202-1012), according to the manufacturer’s instructions. Libraries ranging from 250 to 350 bp were extracted from gel incisions using the QIAquick Gel Extraction Kit (QIAGEN, 28706), PCR amplified, and purified using AMPure XP beads (Beckman Coulter Life Science, A63882). The quality of the resulting libraries was analyzed on the Standard Sensitivity NGS Fragment Analysis Kit (Agilent, DNF-473-0500) on a 12-Capillary Fragment Analyzer. Libraries were quantified using the Quant-iT dsDNA assay kit (Q33120), and equal concentrations (2 nM) of libraries were pooled and subjected to single-end (1 × 150) sequencing of ∼60 to 80 million reads per sample using the High Output v2 kit (Illumina, FC-404-2002) on a NextSeq550 following the manufacturer’s instructions.

#### ChIP-seq analysis

For ChIP-seq, we used Bowtie2 to map reads to the mm10 or hg38 genome^95^. We marked duplicates using the Picard *MarkDuplicates* function and removed duplicates using *‘samtools view -b -F 1024’*. Next, we used deepTools *bamCoverage* to generate RPKM-normalized bigwig tracks and *bamCompare* to compare ChIP to input^96^. We called peaks using the broad mode of MUSIC^97^.

#### ChIP-seq repeat enrichment analysis

We downloaded the repeat regions using the UCSC Genome Browser Table Browser. We selected the RepeatMasker track from the UCSC Table Browser, specifying the appropriate genome build (mm10 or hg38). This dataset was used to perform the enrichment of different repeat elements in our ChIP-seq data. For each repeat element, we calculated an enrichment z-score using in-house scripts. We first stratified the repeat regions into chromosomes. On each chromosome, we moved every repeat element to a random location on the chromosome independently from other repeat regions. After all repeat regions were shuffled, we calculated the nucleotide overlap between the shuffled repeat region set and the PDGFB peaks. The total number of overlapping nucleotides between repeats and PDGFB peaks is used as our enrichment test statistic. The shuffling is repeated 20 times to build an estimate of the null distribution of the overlap between PDGFB and repeat regions. The mean and standard deviation of the expected random overlap distribution was calculated. We finally calculated the z-score by subtracting the mean from the observed number of overlapping nucleotides in PDGFB-repeat regions and normalizing by the standard deviation. Z-score for each repeat element was recorded.

#### ChIP-seq ENCODE enrichment analysis

ENCODE ChIP-Seq peak files (in narrowPeak format) for human (hg38) and mouse (mm10) histone modifications were downloaded from The ENCODE Project portal (https://www.encodeproject.org/)^98^. We selected processed ChIP-Seq narrowPeak files on the portal and downloaded the file list. Each file in the list was downloaded from the portal (accessed September 2023). We found 13,309 peak files for human and 7,810 peak files for mouse. For each peak file, we calculated an enrichment z-score using in-house scripts. We first stratified the peaks into chromosomes. On each chromosome, we moved every peak to a random location on the chromosome independently from other peaks. After all peaks were shuffled, we calculated the nucleotide overlap between the shuffled histone peak set and the PDGFB peaks. The total number of overlapping nucleotides between histone and PDGFB peaks is used as our enrichment test statistic. The shuffling is repeated 20 times to build an estimate of the null distribution of the overlap between PDGFB and histone peaks. The mean and standard deviation of the expected random overlap distribution was calculated. We finally calculated the z-score by subtracting the mean from the observed number of overlapping nucleotides in PDGFB-histone peaks and normalizing by the standard deviation. Z-score for each narrowPeak file was recorded.

#### Single cell RNA-sequencing (scRNA-seq)

Human tumors were prepared as single-cell suspensions. Briefly, samples were coarsely chopped with surgical scissors and enzymatically digested with Papain supplemented with DNase I (Worthington Biochemical Corporation, LK003150). Samples were incubated for 15 min at 37°C on a thermocycler kept at 1400×*g* and briefly pipetted every 5 min. Cells were pelleted at 13,000×*g* for 10 s and resuspended in phosphate-buffered saline (PBS) before processing for debris and dead cell removal. Cell suspensions were processed using the MACS Debris Removal Kit (Miltenyl, 130-109-398) and MACS Dead Cell Removal Kit (Miltenyl, 130-090-101), according to the manufacturer’s instructions. Live cells were collected through negative selection using an MS Column in the magnetic field of a MiniMACS Separator (Miltenyl, 130-042-102). Eluted cells were spun at 300×*g* for 5 min and resuspended in Gibco Dulbecco’s Modified Eagle Medium with GlutaMAX (DMEM; Thermofisher, 10566016) supplemented with 10% fetal bovine serum (FBS; Thermofisher, 16000044). Single cells were processed with the 10X Chromium 3′ Single-Cell Platform using the Chromium Single-Cell 3′ Library, Gel Bead, and Chip Kits (10X Genomics) following the manufacturer’s protocol. Briefly, approximately 5,000-15,000 cells were added to each channel of a chip to be partitioned into Gel Beads in Emulsion (GEMs) in the Chromium instrument, followed by cell lysis and barcoded reverse transcription of RNA in droplets. GEMs were broken, and cDNA from each single cell was pooled. Clean-up was performed using Dynabeads MyOne Silane Beads (Thermofisher, 37002D). Subsequently, the cDNA was amplified and fragmented to optimal size before end repair, A-tailing, and adaptor ligation. Libraries were run individually using a NextSeq 500/550 High Output Kit v2.5 (75 Cycles) (Illumina, 20024907) and sequenced on an Illumina NextSeq550 instrument.

#### Single cell ATAC-sequencing (scATAC-seq)

Single cell ATAC Library was prepared according to Chromium Next GEM Single cell ATAC Reagent kit v2 (10x Genomics). In Brief, prepared nuclei were incubated with transposome and transposase entered and preferentially fragmented DNA in open region of chromatin. Transposed single nuclei, a master mix, Gel Beads containing barcoded oligonucleotides, and oil were loaded on a Chromium controller (10x Genomics) to generate GEMS (Gel Beads-In-Emulsions) where barcoded single strand DNA was synthesized. Subsequently the GEMS are broken and pooled. Following sequential cleanup using Dynabeads MyOne Silane Beads and SPRI beads, barcoded DNA fragments are amplified by PCR to generate indexed library.

#### scRNA-seq and scATAC-seq processing and analysis

Samples for scRNA-seq and scATAC-seq were run on a 10X Genomics Chromium platform to produce next-generation sequencing libraries. Reads were aligned to the mouse genome (mm10) and human genome (hg38) using Cell Ranger. For scRNA-seq, standard procedures for filtering, mitochondrial gene removal, and variable gene selection were used using the Seurat pipeline^99^. The criteria for cell and gene inclusion in human and mouse scRNA-seq were as follows: genes present in more than three cells were included, cells that expressed >300 genes were included, the number of genes detected in each cell was >200 and <5000, and the mitochondria ratio was less than 10. We integrated cells from different patients and experiments using the Harmony algorithm^100^. Next, we visualized clusters using a uniform manifold approximation and projection constructed from the Harmony-corrected PCA. This visualization was created using the *runUMAP*, *FindNeighbors*, and *FindClusters* functions of the Seurat package. We extracted differentially expressed genes among clusters using *FindAllMarkers* function of Seurat package.

We performed enrichment analysis using enrichR R package^101^. Next, we used well-established cell type markers to annotate each cluster with its corresponding cell type and also ran SCRAM for annotating each cell in single-cell resolution^41,102–104^.

For scATAC-seq, we first created a unified set of peaks using functions from the GenomicRanges package, merging all intersecting peaks with the reduce function. To quantify our combined set of peaks, we created a Fragment object for each experiment. We quantified counts in peaks using the FeatureMatrix function in Signac^105^. We ran NucleosomeSignal and TSSEnrichment for quality control. For dimension reduction, we applied FindTopFeatures, RunTFIDF, RunSVD, and RunUMAP. Cell clustering was performed using FindNeighbors and FindClusters, reducing dimensionality by applying LSI and UMAP with LSI components 2 to 30. We inferred gene activities using the GeneActivity function in Signac. To add the DNA sequence motif information required for motif analyses, we used the AddMotifs function in Signac. To identify potentially important cell-type-specific regulatory sequences, we searched for DNA motifs overrepresented in differentially accessible peaks between cell types.

#### AlphaFold and protein interaction prediction analysis

We used AlphaFold2^106,107^ to predict the protein structure of the full preprotein form of PDGFB and the C-terminal containing forms of PDGFB^wt^ and PDGFB^ΔNLS^ showing conformational changes to the disordered C-terminus upon neutralizing K and R residue using full databases with default parameters.

## Statistical analysis

Sample sizes and statistical tests are provided in the figure legends. The following tests were used for statistical analysis, unless otherwise noted. For Kaplan-Meier survival analysis, the log-rank test was used to compare survival differences across groups. For statistical analyses, a two-tailed Student’s t-test was used unless noted otherwise in the corresponding figure legend. Significant differences are denoted by asterisks in associated graphs. Data are presented as the mean±standard error of the mean. Levels of statistical significance are indicated as follows: ns: not significant, \**p*<0.05, \*\**p*<0.01, \*\*\**p*<0.001, and \*\*\*\**p*<0.0001.

## Supplemental Figures and Tables

**Figure S1.**
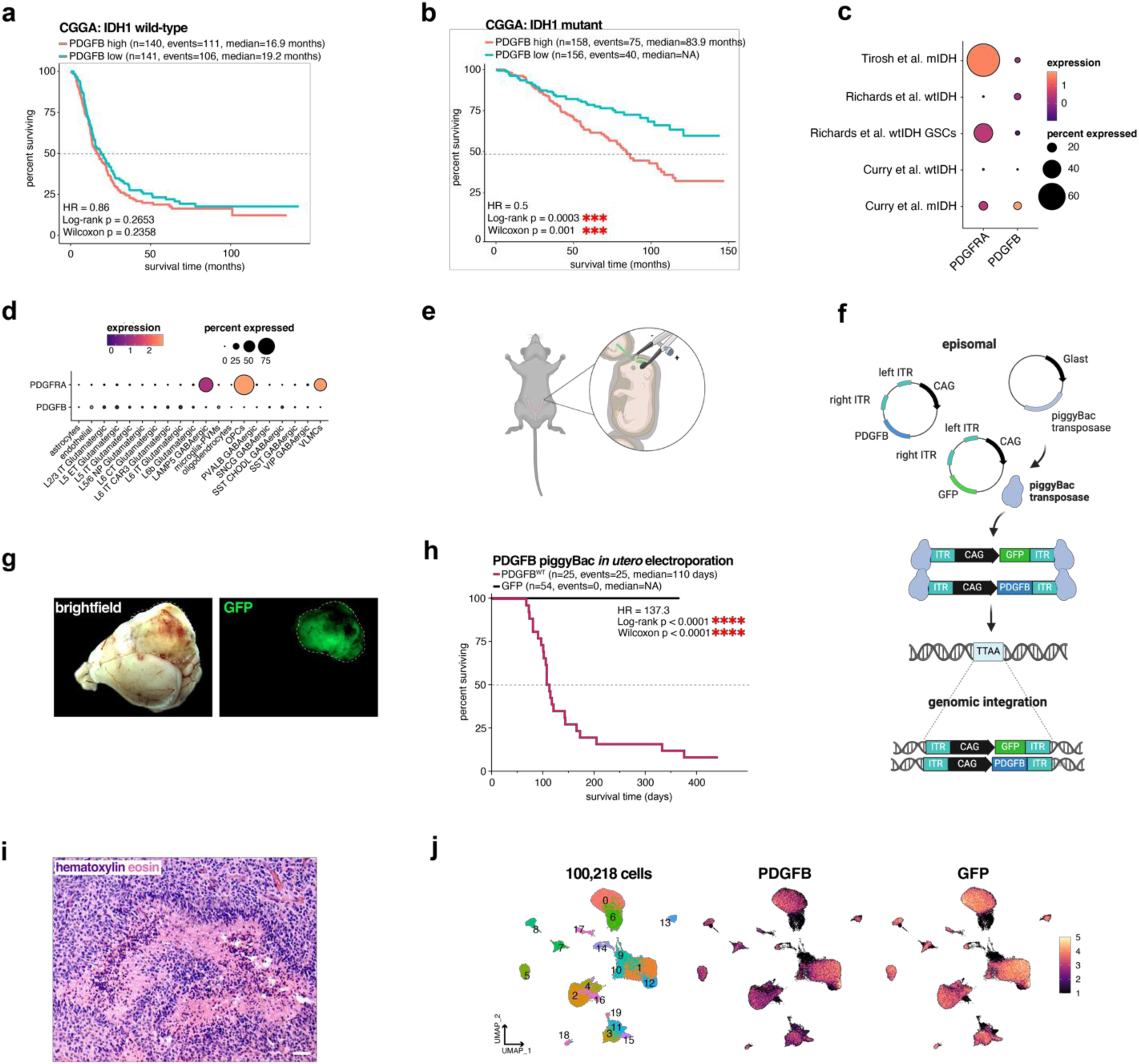
PDGFB PDGFB drives mouse gliomagenesis in the absence of mutations. (a) Kaplan-Meier survival analysis of wtIDH glioma patients from the CGGA split on median *PDGFB* expression. No significant difference in survival outcome is observed between high (n=140; events=111; median=16.9 months) and low (n=141; events=106; median=19.2 months) *PDGFB* expression groups (HR=0.86; log-rank *p*<0.2653; Wilcoxon *p*<0.2358). (b) Kaplan-Meier survival analysis of mtIDH glioma patients from the CGGA split on median *PDGFB* expression. High *PDGFB* expression (n=158; events=75; median=83.9 months) is associated with worse survival outcomes when compared to low *PDGFB* expression (n=158; events=40; median=NA) (HR=0.5; log-rank \*\*\**p*<0.0003; Wilcoxon *\*\*\*p*<0.001). (c) Dot plot of scRNA-seq data from previously published datasets showing *PDGFRA* and *PDGFB* expression are higher in mIDH glioma as compared to wtIDH glioma. (d) Dot plot of non-tumor human brain scRNA-seq data from the Allen Brain Atlas shows *PDGFB* is minimally expressed in the non-tumor human brain and *PDGFRA* is enriched in OPCs. (e) Schematic representation of the intraventricular injection used in the piggyBac *in utero* electroporation system. (f) Schematic representation of the expression vectors used in the piggyBac *in utero* electroporation system. (g) Brightfield and fluorescent images of whole brains showing end-stage tumors in PDGFB^wt^ mice; white dashed line denotes tumor. (h) Kaplan-Meier survival analysis of PDGFB^wt^ (n=25; events=25; median=110 days) and GFP alone (n=54; events=0; median=NA) mice (HR=137.3; log-rank \*\*\*\**p*<0.0001; Wilcoxon *\*\*\*\*p*<0.0001). (i) H&E staining of an end-stage PDGFB^wt^ (P100) tumors shows histopathology consistent with high-grade glioma; scale bar = 50 microns. (j) UMAPs of scRNA-seq (n=100,218 cells; n=3 mice) data shows GFP and PDGFB overexpression transcripts co-localize to the same cells. CGGA: Chinese Glioma Genome Atlas; H&E: hematoxylin and eosin.

**Figure S2.**
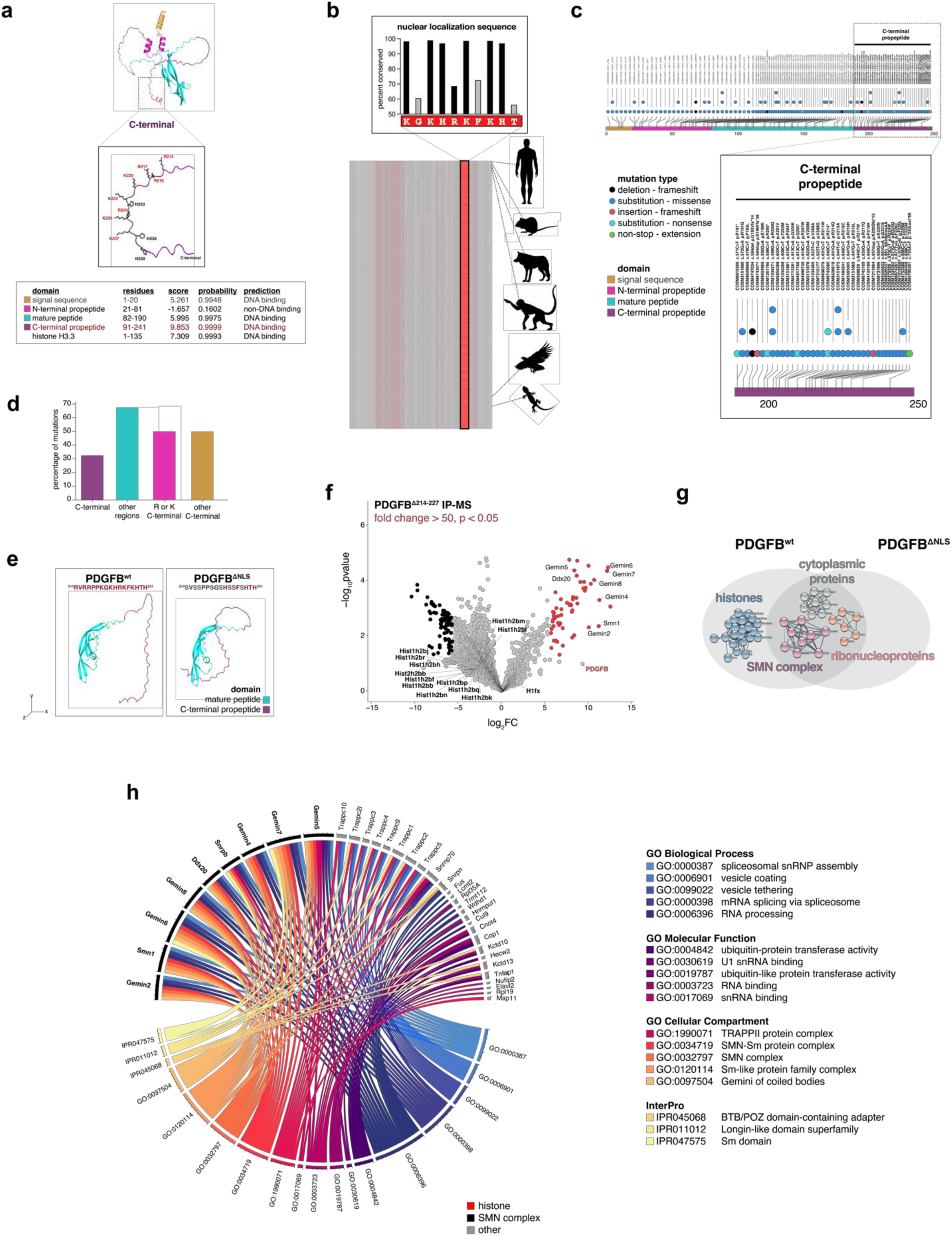
PDGFB contains a highly conserved nuclear localization sequence at its C– terminus. (a) AlphaFold2 modeling of pre-PDGFB. Inset showing basic amino acid residues at the C-terminus and Schematic representation of PDGFB protein domains showing basic amino acid residue enrichment in at the C-terminus. Underlined sequence denotes NLS. Putative DNA-binding prediction scores are shown by protein domain. Histone H3.3 score is shown for reference. (b) Snapshot of PDGFB ortholog alignment across 600 vertebrate genomes shows NLS sequence is highly conserved at most K and R residues. (c–d) Lollipop plot and quantification of TCGA pan-cancer datasets showing the highest incidence of mutations (number of mutations/number of amino acids in each protein domain) is found in the C-terminal domain of PDGFB. (e) AlphaFold2 modeling of the C-terminal containing forms of PDGFB^wt^ and PDGFB^ΔNLS^ showing conformational changes to the disordered C-terminus upon neutralizing K and R residues. (f) Volcano plot of the 59 proteins identified from PDGFB IP-MS of PDGFB^ΔNLS^ tumors. Red dots denote proteins with fold change>50; *p*<0.05; black dots denote proteins enriched in IgG fold change<50; *p*<0.05. (g) STRING network of significant proteins from *(f)* showing histone enrichment is no longer detected and schematic representation of significant proteins found in PDGFB^wt^ and PDGFB^ΔNLS^ tumor brains shows histone binding is only detected in PDGFB^wt^ tumors. (h) Circos plot of GOs (*p*<0.05) corresponding to significant proteins from *(f).* NLS: nuclear localization sequence.

**Figure S3.**
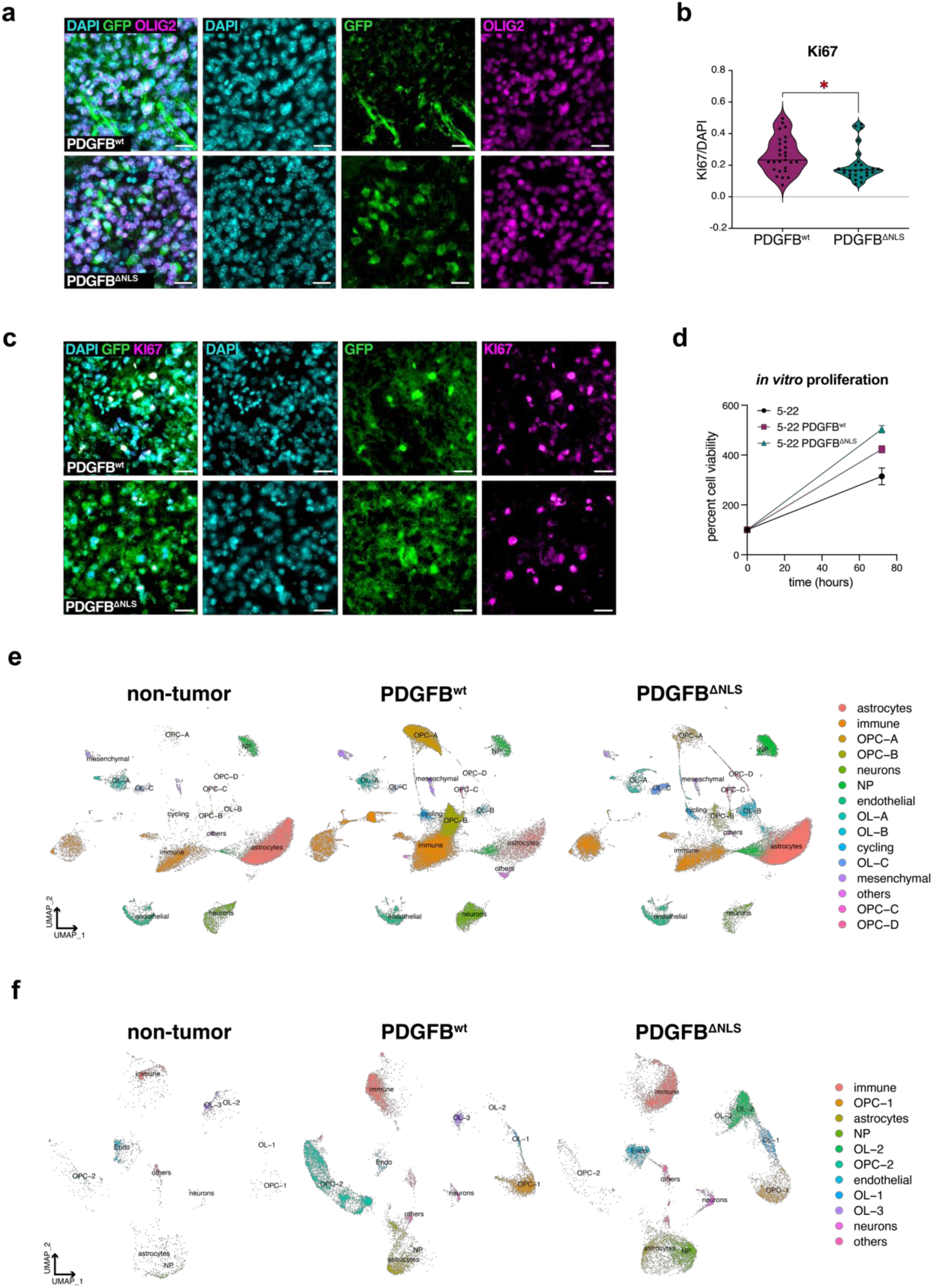
PDGFB drives expansion of OPCs and nuclear PDGFB is required to prevent OL differentiation. (a) Immunostaining of PDGFB^wt^ and PDGFB^ΔNLS^ tumors shows tumor cells are Olig2+; scale bar = 20 microns. (b) Quantification of Ki67-positivity shows both PDGFB^wt^ and PDGFB^ΔNLS^ tumors are highly proliferative, with PDGFB^wt^ tumors showing a higher rate of proliferation. (c) Ki67 immunostaining of PDGFB^wt^ and PDGFB^ΔNLS^ tumors; scale bar = 20 microns. (d) *In vitro* viability assay of mIDH GSCs (5-22) overexpressing PDGFB^wt^ and PDGFB^ΔNLS^ show increased proliferation as compared to controls. (e) Dim plots of 98,242 cells from our scRNA-seq dataset from PDGFB^wt^ (n=3), PDGFB^ΔNLS^ (n=3) and non-tumor (n=3) mouse brains showing cell type annotated clusters. (f) Dim plot of 27,836 cells from our scATAC-seq dataset from PDGFB^wt^ (n=1), PDGFB^ΔNLS^ (n=1) and non-tumor (n=1) mouse brains showing cell type annotated clusters. GSC: glioma stem cell.

**Figure S4.**
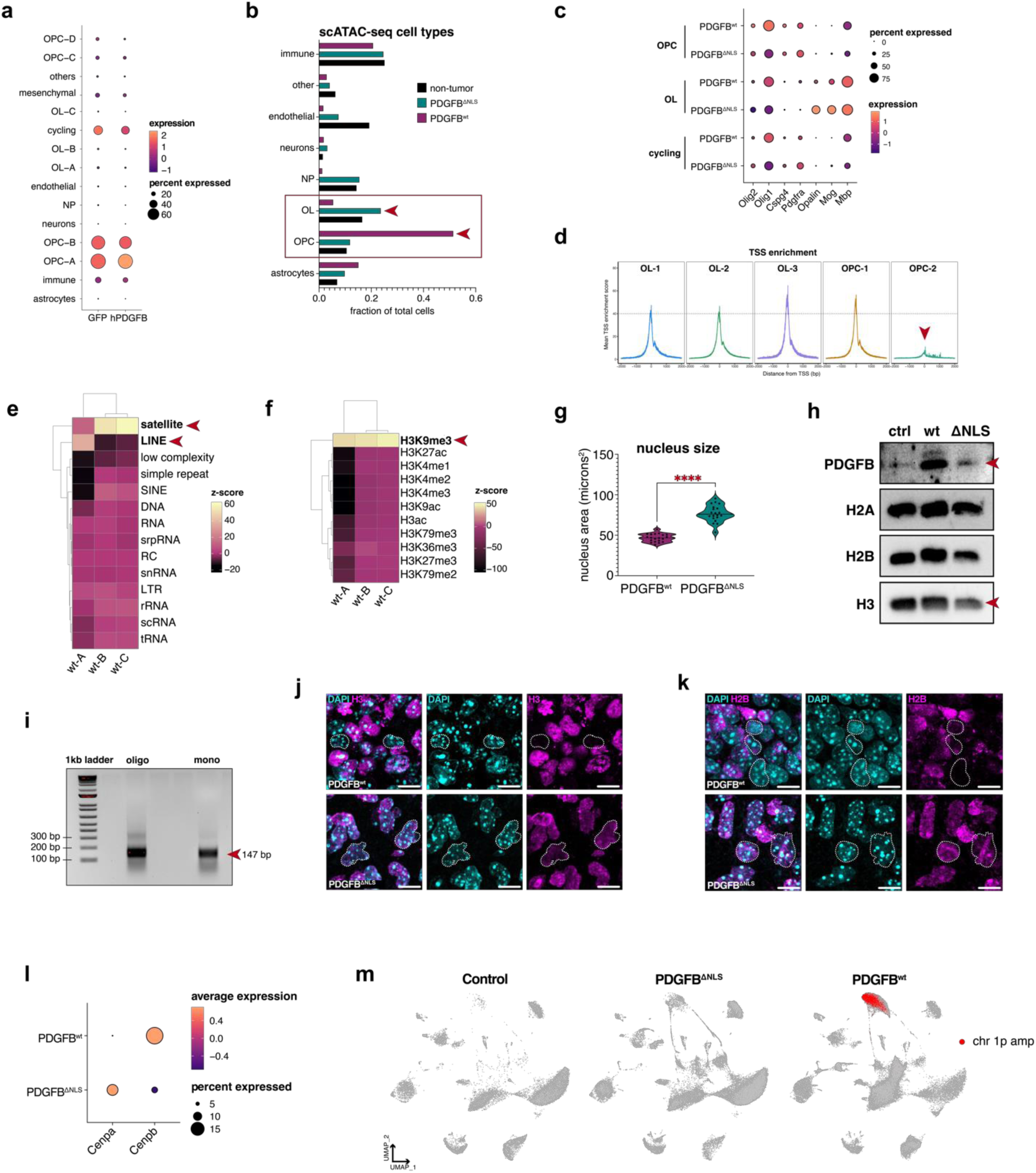
Nuclear PDGFB regulates OPC lineage and heterochromatin architecture. (a) Dot plot of scRNA-seq dataset from PDGFB^wt^, PDGFB^ΔNLS^ and non-tumor mouse brains shows OPCs and cycling tumor cells express *GFP* and *PDGFB*. (b) Bar plot of OPC and OL lineage clusters from scATAC-seq showing the OPCs are expanded in PDGFB^wt^ tumors whereas OLs are increased in PDGFB^ΔNLS^ tumors. (c) Dot plot of OPC, OL and cycling tumor cells from scRNA-seq dataset shows OPCs are enriched for OPC lineage markers in PDGFB^wt^ tumors and OLs are enriched for OL lineage markers in PDGFB^ΔNLS^ tumors. (d) TSS analysis from scATAC-seq dataset shows OPC-2 cells, which are predominantly found in PDGFB^wt^ tumors, show a large reduction chromatin accessibility. (e–f) Enrichment analysis of PDGFB ChIP-seq from mouse PDGFB^wt^ tumors shows enrichment in PDGFB binding to *(e)* satellite DNA regions (e.g. centromeric repeats), long interspersed nuclear elements (LINEs) and *(f)* H3K9me. (g) Quanitification of nucleus size shows PDGFB^wt^ nuclei are smaller than PDGFB^ΔNLS^ nuclei. (h) Immunoblot of nucleosome extractions shows PDGFB is detected in the nucleosome fraction and reduced H3 levels in both PDGFB^wt^-and PDGFB^ΔNLS^-overexpression wtIDH GSCs (7-2). (i) Image of agarose gel showing single nucleosomes were obtains for the nucleosome extraction immunoblot performed in *(h)*. (j–k) Representative images from *(j)* H3 and *(k)* H2B immunostaining from PDGFB^wt^ and PDGFB^ΔNLS^ tumors. White dashed lines show select cells with reduced H3 and H2B positivity in PDGFB^wt^ tumors and nuclei with aberrant morphology in PDGFB^ΔNLS^ tumors. (l) Dot plot from PDGFB^wt^ and PDGFB^ΔNLS^ mouse tumor scRNA-seq dataset shows *Cenpb* expression is increased in PDGFB^wt^ tumor cells and is reduced when PDGFB cannot enter the nucleus. (m) Feature plots showing scRNA-seq inferred chromosome 1p amplifications are present in PDGFB^wt^ tumors.

**Figure S5.**
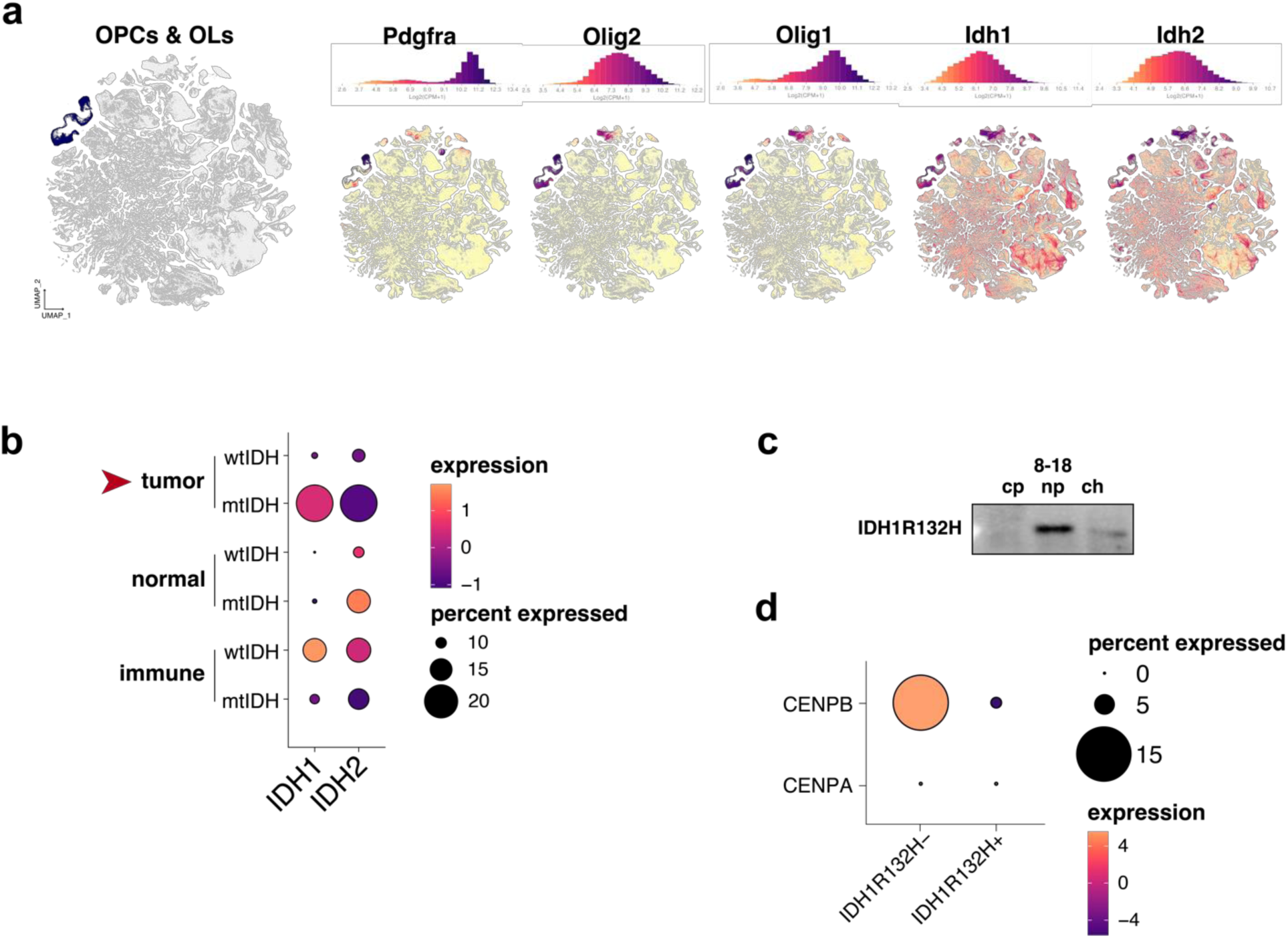
OPCs and mIDH glioma cells are enriched for *IDH1* and *IDH2*. (a) Dim plot and corresponding feature plots are shown for 4.04 million cells from Allen Brain Cell Atlas. Feature plots for OPC and OL lineage markers show *Idh1* and *Idh2* are highly expressed in OPC and OL cells in normal mouse brain. (b) Dot plot of human brain scRNA-seq data from Allen Brain Atlas shows *IDH1* expression is highest in OPCs. (c) Immunoblot of cell fractions from mIDH GSCs (8-18) shows endogenous IDH1^R132H^ is detected in the nucleoplasm. (d) Dot plot from human glioma scRNA-seq dataset showing IDH1^R132H^– cells have increased *CENPB* expression as compared to IDH1^R132H^+ cells.

**Table S1.** Characteristics of human glioma samples used for this study.

| Patient | Tumor Type | IDH Status | 1p19q codeletion | WHO Grade | Recurrence | Experiments |
| --- | --- | --- | --- | --- | --- | --- |
| WT-01 | astrocytoma (GBM) | wild-type | no | IV | primary | ChIP |
| WT-02 | astrocytoma (GBM) | wild-type | no | IV | primary | ChIP |
| WT-03 | astrocytoma (GBM) | wild-type | no | IV | primary | ChIP |
| WT-04 | astrocytoma (GBM) | wild-type | no | IV | primary | ChIP |
| WT-05 | astrocytoma (GBM) | wild-type | no | IV | primary | ChIP |
| A-01 | astrocytoma | mutant | no | II | primary | ChIP |
| O-01 | oligodendroglioma | mutant | yes | II | primary | ChIP |
| O-02 | oligodendroglioma | mutant | yes | II | recurrent | ChIP; ChIP-seq |
| A-02 | astrocytoma | mutant | no | IV | primary | ChIP; ChIP-seq |
| O-03 | oligodendroglioma | mutant | yes | II | primary | ChIP; ChIP-seq |
| O-04 | oligodendroglioma | mutant | yes | III | primary | ChIP |
| A-03 | astrocytoma | mutant | no | III | primary | ChIP; IHC |
| A-04 | astrocytoma | mutant | no | II | primary | IHC |
| A-05 | astrocytoma | mutant | no | IV | primary | IHC |

